# Hydroxylated monoterpenes mimic bacterial attack to trigger jasmonate-dependent self-amplifying immunity in tomato

**DOI:** 10.64898/2026.06.09.731038

**Authors:** Julia Pérez-Pérez, Pavel Brito-Gutiérrez, Antonio Santiago, Maite Sanmartín, José Tomás Matus, Maria Sulli, Gianfranco Diretto, Francisco Vera-Sirera, Ismael Rodrigo, María Pilar López-Gresa, Purificación Lisón

## Abstract

Hydroxylated monoterpenes (HMTPs) are emitted during resistant tomato-*Pseudomonas syringae* interactions and confer antibacterial resistance, yet their integration into immune signalling remains poorly understood. Here we show that HMTPs act as endogenous mimics of pathogen attack that engage canonical defence pathways in tomato. Using α-terpineol as a representative HMTP, we demonstrate that this volatile activates MPK kinase-, calcium- and reactive oxygen species (ROS)-dependent signalling, promotes jasmonate and salicylic acid accumulation, and induces pathogen-like stomatal immunity independently of abscisic acid. Functional analyses revealed that HMTPs biosynthesis depends on ROS and jasmonate signalling, and HMTPs further promote their own accumulation, establishing a self-reinforcing feed-forward mechanism. Moreover, *in vitro* oxidative conditions drive chemical remodelling and selective interconversion among HMTPs, contributing to volatile diversification and favouring the accumulation of highly bioactive hydroxylated forms. Consistently, deuterium-labelled linalool is incorporated into plant metabolism and converted into deuterated α-terpineol *in planta*, providing direct evidence for volatile interconversion. Together, our findings establish HMTPs as dynamic amplifiers of tomato antibacterial immunity and key actors in pathogen-associated signalling.

## INTRODUCTION

The continuous interaction between plants and their environment has driven the evolution of sophisticated defence mechanisms through intricate signalling networks. Early immune responses involve the rapid production of reactive oxygen species (ROS), transient increases in cytosolic calcium (Ca^2+^) levels, and activation of mitogen-activated protein kinase (MPK) cascades, which integrate early stress perception with downstream transcriptional and metabolic reprogramming (Ali et al., 2024). It is well-known that salicylic acid (SA) and jasmonic acid (JA) pathways are central regulators of plant immunity. SA acts as a key signal in systemic acquired resistance (SAR), modulating local and systemic defences through finely controlled modifications such as glycosylation and methylation (Wildermuth et al., 2021; Bellés et al., 2006; Ding et al., 2020). Conversely, JA and related oxylipins govern stress adaptation, controlling transcriptional reprogramming in response to wounding and pathogen attack (Ruan et al., 2019). JA biosynthesis involves sequential oxidation of α-linolenic acid by lipoxygenases (LOXs), and conversion via allene oxide synthase (AOS) and allene oxide cyclase (AOC) to produce 12-oxo-phytodienoic acid (OPDA), further processed through β-oxidation to yield JA which, upon conjugation to isoleucine, generates the bioactive signal JA-Ile (Li et al., 2021).

While the classical view considers JA an antagonist of SA-mediated defences against biotrophic pathogens, rapid local and systemic JAs signalling is essential for the initiation and establishment of SAR following effector-triggered immunity (ETI), revealing a central role for JAs in long-distance immune signalling (Gaikwad et al., 2026). Beyond hormonal pathways, plant immunity relies on metabolic crosstalk, where SA production and volatile terpene biosynthesis are metabolically interconnected through the plastidial methyl-erythritol phosphate (MEP) pathway (Xiao et al., 2021). Terpenes function both as direct antimicrobials and as volatile organic compounds (VOCs) that modulate plant defence responses (Brilli et al., 2019). Terpenoids are produced via the mevalonic acid (MVA) and MEP pathways, resulting in monoterpenes, sesquiterpenes, and diterpenes (Degenhardt et al., 2009), which can undergo oxidative modifications catalysed by cytochrome P450 monooxygenases (CYPs) to form hydroxylated monoterpenes (HMTPs) (Pichersky et al., 2018).

In tomato (*Solanum lycopersicum*), the terpene synthase (*TPS*) gene family comprises 34 functional genes encoding enzymes with distinct substrate specificities (Zhou et al., 2020). *TPS* gene expression is regulated by multiple stimuli, including herbivory, wounding, and JA signalling (Howe et al., 2008; Dudareva et al., 2013), mimicking herbivore-induced responses across angiosperms (Arimura et al., 2004; Gomez et al., 2005). While terpenoids have traditionally been studied for defence against herbivores, their involvement in plant immunity against microbial pathogens is increasingly recognized. In tomato, HMTPs such as α-terpineol, linalool, and terpinen-4-ol are differentially emitted upon bacterial recognition triggering ETI, and their production is linked to the induction of the monoterpene synthase gene *SlMTS1* (van Schie et al., 2007; López-Gresa et al., 2027).

However, the biochemical origin of key HMTPs such as α-terpineol remains unresolved, as no terpineol synthase activity has been described within the tomato *TPS* family ^12^. Despite this discrepancy, HMTPs actively contribute to defence against the bacterial pathogen *Pseudomonas syringae* pv. tomato (Pst) (Riedlmeier et al., 2017; Wenig et al., 2019; Vlot et al., 2021; Pérez-Pérez et al., 2024). In tomato, alterations in HMTP production impact SA accumulation, revealing a tight metabolic coordination between terpene-derived VOC biosynthesis and SA-mediated immunity (Pérez-Pérez et al., 2024). Functional assays revealed that these compounds induce stomatal closure, with α-terpineol displaying the strongest phenotypic effects and disease resistance (Pérez-Pérez et al., 2024). Nevertheless, the specific mode of action of α-terpineol and its integration into defence signalling pathways remain largely unexplored. Here, by integrating transcriptomics, metabolomics, and biochemical analyses, we demonstrate that α-terpineol activates defence-associated signalling and coordinates its activity with key phytohormones. Moreover, we uncover a self-reinforcing regulatory process controlling HMTP levels during ETI establishment and reveal a novel chemical, non-enzymatic remodelling process underlying sustained volatile diversification during plant immune responses.

## RESULTS

### α-Terpineol elicits a transcriptional reprogramming resembling *Pseudomonas syringae* infection

We have previously shown that HMTPs levels, including α-terpineol, linalool and terpinen-4-ol, increase during ETI in tomato upon infection with *Pseudomonas syringae*, being α- terpineol the most effective HMTP in promoting stomatal closure, suggesting a predominant role in plant defence (Pérez-Pérez et al., 2024). Based on these observations, we hypothesized that α-terpineol could partially mimic defence responses triggered during Pst infection, thereby conferring resistance through activation of transcriptional defence mechanisms. To test this, we performed RNA-seq analysis under four experimental conditions: α-terpineol-treated and untreated control tomato plants, as well as Pst-infected and mock plants.

α-Terpineol treatment triggered a markedly broader transcriptional reprogramming (5,809 differentially expressed genes) compared to the more moderate response induced by Pst infection (2,742 genes) (Fig. 1A). This suggests that α-terpineol may exert broader effects beyond its defensive role. Despite these differences, α-terpineol-induced transcriptional changes overlapped with 44.6% of the genes responsive to Pst infection, highlighting a shared transcriptional signature. This relationship is supported by a highly significant correlation (p < 2.2 × 10⁻¹⁶; Pearson’s r = 0.74), indicating a strong correspondence between the transcriptional activities induced by α-terpineol and those triggered by bacterial infection (Fig. 1B). Volcano plot analyses further illustrate this predominant transcriptional activation response (Supplementary information, Fig. S1A). α-Terpineol treatment induced the expression of the immune marker genes, as determined by RT–qPCR (Supplementary information, Fig. S1B), confirming FPKM analysis and the reliability of the RNA-seq data (Supplementary information, Fig. S1C).

**Figure 1.**
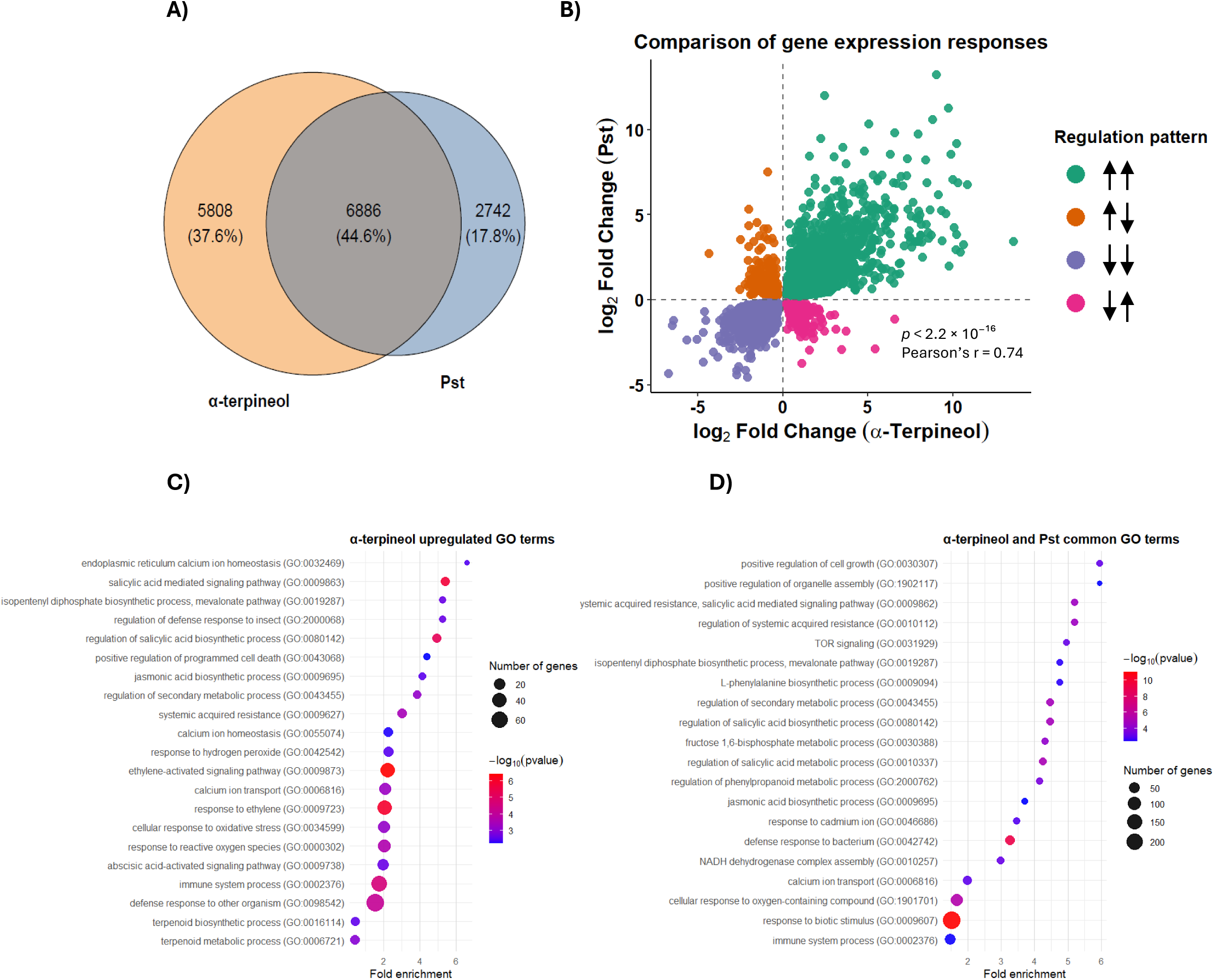
Transcriptional comparison between α-terpineol treatment and *Pseudomonas syringae* infection. **(A)** Venn diagram showing the number of differentially expressed genes (DEGs) identified in plants treated with α-terpineol and in plants infected Pst (*P. syringae*), each compared to their respective untreated controls. Numbers indicate the total DEGs unique to each condition and those shared between both treatments**. (B)** Scatter plot showing the log₂ fold change (log₂FC) of DEGs in response to α-terpineol treatment (x-axis) and *P. syringae* infection (y-axis). Each dot represents one gene, and colours indicate the four quadrants corresponding to the direction of regulation in each condition. The dashed line represents the diagonal (y = x). The first arrow corresponds to the Pst data, and the second arrow corresponds to the α-terpineol. Gene Ontology (GO) enrichment analysis of biological process terms associated with genes **(C)** upregulated and **(D)** common GO terms regulated by α-terpineol treatment and Pst infection, are displayed as bubble plot, showing enriched Gene Ontology (GO) biological processes associated with α-terpineol and Pst treatment. The x-axis represents fold enrichment, bubble size indicates the number of genes associated with each GO term, and colour intensity corresponds to statistical significance as −log10(*p*-value).

Gene ontology (GO) enrichment analysis of α-terpineol-dependent genes (Fig. 1C) showed a massive enrichment of core immunity-related terms, most notably the ‘salicylic acid mediated signalling pathway’, ‘defence response to other organism’, ‘systemic acquired resistance (SAR)’, ‘jasmonic acid biosynthetic process’ and ‘response to ethylene’, revealing a multi-layered immune state. Furthermore, the strong enrichment of ‘calcium ion homeostasis’ suggests that this compound may trigger essential signalling events typically attributed to pathogen recognition. Specialized metabolism is also heavily prioritized, with significant enrichment in the ‘terpenoid biosynthetic process’ and the ‘mevalonate pathway’, indicating that α-terpineol may induce a metabolic ‘priming’ effect. Shared GO terms between α-terpineol and Pst (Fig. 1D), such as ‘TOR signalling’ and ‘positive regulation of cell growth’, highlight a sophisticated coordination between growth and immunity.

### α-Terpineol induces pathogen-like stomata closure by acting on core defence signalling pathways

To determine whether the signalling pathways underlying HMTPs-mediated stomata closure overlap with those activated during bacterial infection, we conducted stomatal aperture assays using tomato leaf discs and specific pharmacological inhibitors (Fig. 2). To investigate Ca^2+^ signalling, we pre-treated samples with the calcium chelator EGTA (ethylene glycol tetraacetic acid); for MPK cascade involvement, we used PD98059, and for ROS production, we applied DPI (diphenyleneiodonium, a NADPH oxidase inhibitor), SHAM (salicylhydroxamic acid, an inhibitor of the mitochondrial alternative oxidase), and NAC (N-acetylcysteine, a potent antioxidant). Our results showed that EGTA completely abolished stomatal closure induced by any treatment, indicating that Ca^2+^ signalling is essential for the induction of stomatal immunity responses (Fig. 2A). Treatment with PD98059 (Fig. 2B) or DPI (Fig. 2C) partially inhibited stomatal closure induced by α-terpineol, flg22, and ABA, suggesting that MPK cascades and NADPH-dependent ROS production are partially required. Conversely, SHAM did not inhibit ABA and α-terpineol-induced stomatal closure (Fig. 2D), while treatment with NAC abolished stomatal closure triggered by both α-terpineol and flg22 (Fig. 2E), revealing a shared requirement for ROS accumulation that distinguishes their mode of action from ABA.

**Figure 2.**
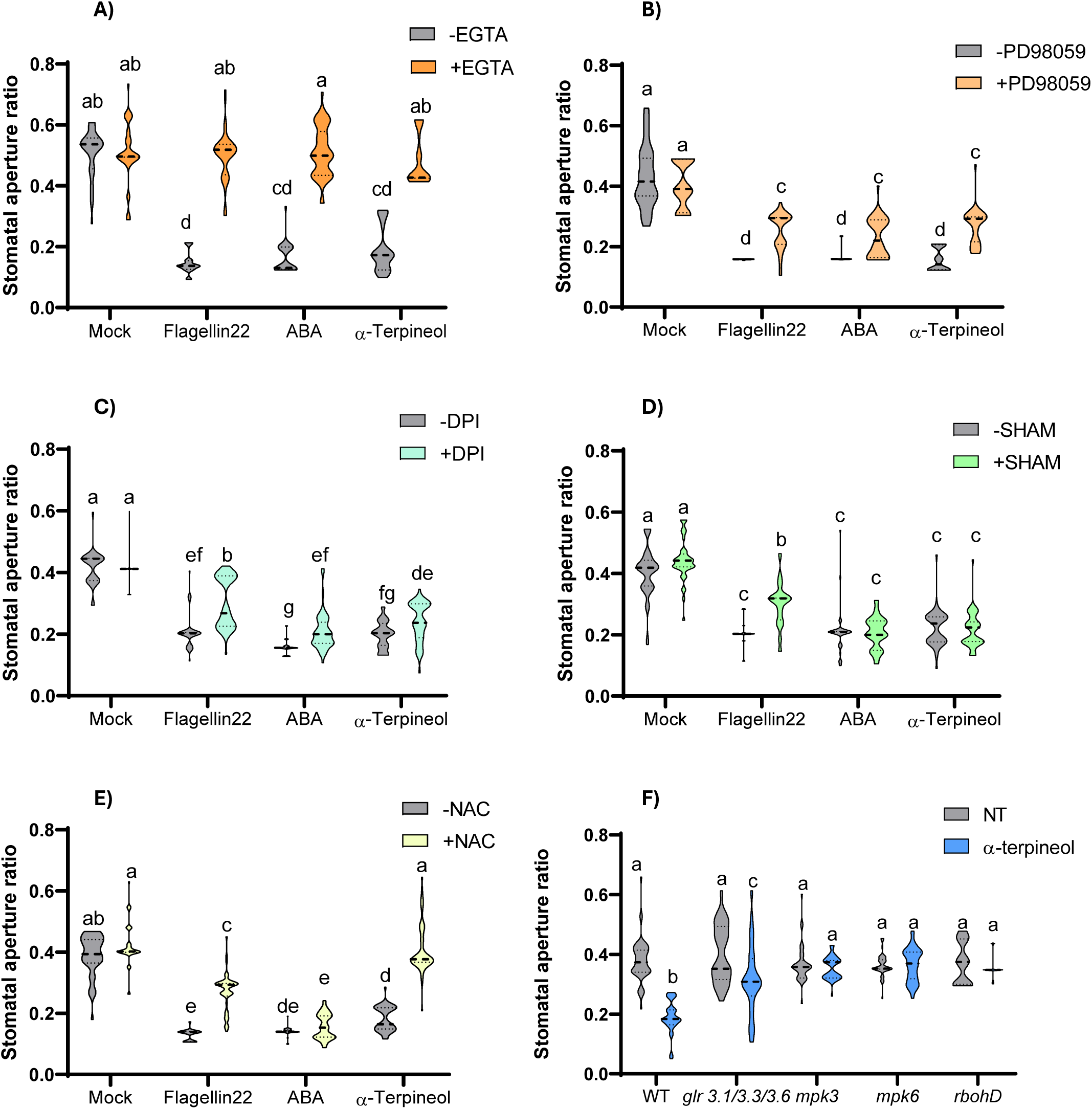
Effect of pharmacological inhibitors on stomatal aperture responses to α-terpineol and control treatments. **(A)** Stomatal aperture ratio measured in epidermal peels treated with mock solution, flagellin22, abscisic acid (ABA), or α-terpineol in the absence (−EGTA) or presence (+EGTA) of the calcium chelator EGTA. **(B)** Stomatal aperture ratio measured after the same treatments in the absence (−PD98059) or presence (+PD98059) of the MPK (Mitogen-Activated Protein Kinases) pathway inhibitor PD98059. **(C)** Stomatal aperture ratio measured in the absence (−DPI) or presence (+DPI) of the NADPH oxidase inhibitor diphenyleneiodonium (DPI). **(D)** Stomatal aperture ratio measured in the absence (−SHAM) or presence (+SHAM) of salicylhydroxamic acid (SHAM). **(E)** Stomatal aperture ratio measured in the absence (−NAC) or presence (+NAC) of the antioxidant N-acetylcysteine (NAC). **(F)** Stomatal aperture ratio (width/length) measured in wild-type (WT), *glr3.1/3.3/3.6*, *mpk3*, *mpk6*, *rbohD* plants under non-treated (NT) conditions or after treatment with α-terpineol. Violin plots show the distribution of stomatal aperture ratios (n=30), with central values indicated for each treatment. Different letters indicate statistically significant differences among treatments (*p* < 0.05).

To genetically validate these observations, we used several Arabidopsis mutants. The triple knock-out *glr* mutant (*glr3.1/3.3./3.6*), which is compromised in Ca^2+^ signals mediated by glutamate receptor-like channels, was partially sensitive to α-terpineol treatment on stomatal closure. Mitogen-activated kinase mutants *mpk3, mpk6,* and NADP oxidase mutants *rbohD* resulted to be completely insensitive to α-terpineol (Fig. 2F). Parallel experiments conducted with linalool revealed an identical behavioural pattern (Supplementary information, Fig. S2A-F), further confirming that both HMTPs trigger stomatal closure through the same core signalling modules but α-terpineol conferred the strongest resistance to Pst among the monoterpenoids tested (Supplementary information, Fig. S2G and H, Pérez-Pérez et al., 2024).

To investigate if α-terpineol-induced stomatal closure is mediated through ABA signalling, we employed an ABA-responsive luciferase reporter Arabidopsis line, *pMAPKKK18-LUC+* (García-Maquilón et al., 2021), in which α-terpineol exposure failed to induce luciferase activity (Supplementary information, Fig. S3A), consistent with unchanged ABA accumulation levels (Supplementary information, Fig. S3B). However, transcriptomic analysis (Supplementary information, Fig. S3C) revealed a significant induction of marker genes typically associated with ABA signalling (González-Guzmán et al., 2014), such as *SlRAB18* and *SlLEA5* which can be induced by ROS (Karpinska et al., 2022; Evans et al., 2024).RT-qPCR confirmed that α-terpineol treatment significantly upregulated MPK-associated markers, (Supplementary information, Fig. S3D), ROS-related responses (peroxidase, SOD, Supplementary information, Fig. S3E), and Ca^2+^-responsive genes (Supplementary information, Fig. S3F) (Hu et al., 2022). Furthermore, inhibition assays combined with bacterial growth measurements showed that both the MPK cascade and NADPH-dependent ROS production are essential for the full resistance phenotype mediated by α-terpineol (Supplementary information, Fig. S4B-D).

To investigate calcium signalling dynamics beyond its transcriptional activation (Fig. 3A), we utilized *Arabidopsis thaliana* plants constitutively expressing the GCaMP3 cytosolic Ca^2+^ sensor (Nguyen et al., 2018; Pantazopoulou et al., 2023) (Fig. 3B). α-Terpineol application triggered a fast and transient increase in the cytosolic Ca^2+^ concentration ([Ca^2+^]cyt) in the treated leaf, which was later transmitted to distal non-treated leaves (Fig. 3B; Supplementary information, Video S1). These [Ca^2+^]cyt transients were first observed close to the vasculature and later diffused to the blade, suggesting long-distance Ca^2+^ signalling transmitted through the vascular network. In the triple knock-out mutant *glr3.1/3.3./3.6*^29^, no changes in [Ca^2+^]cyt fluxes were observed in any of the leaf blades after α-terpineol application, although a very slight increase was detected in the petioles (Fig. 3B, bottom row, Video S2).

**Figure 3.**
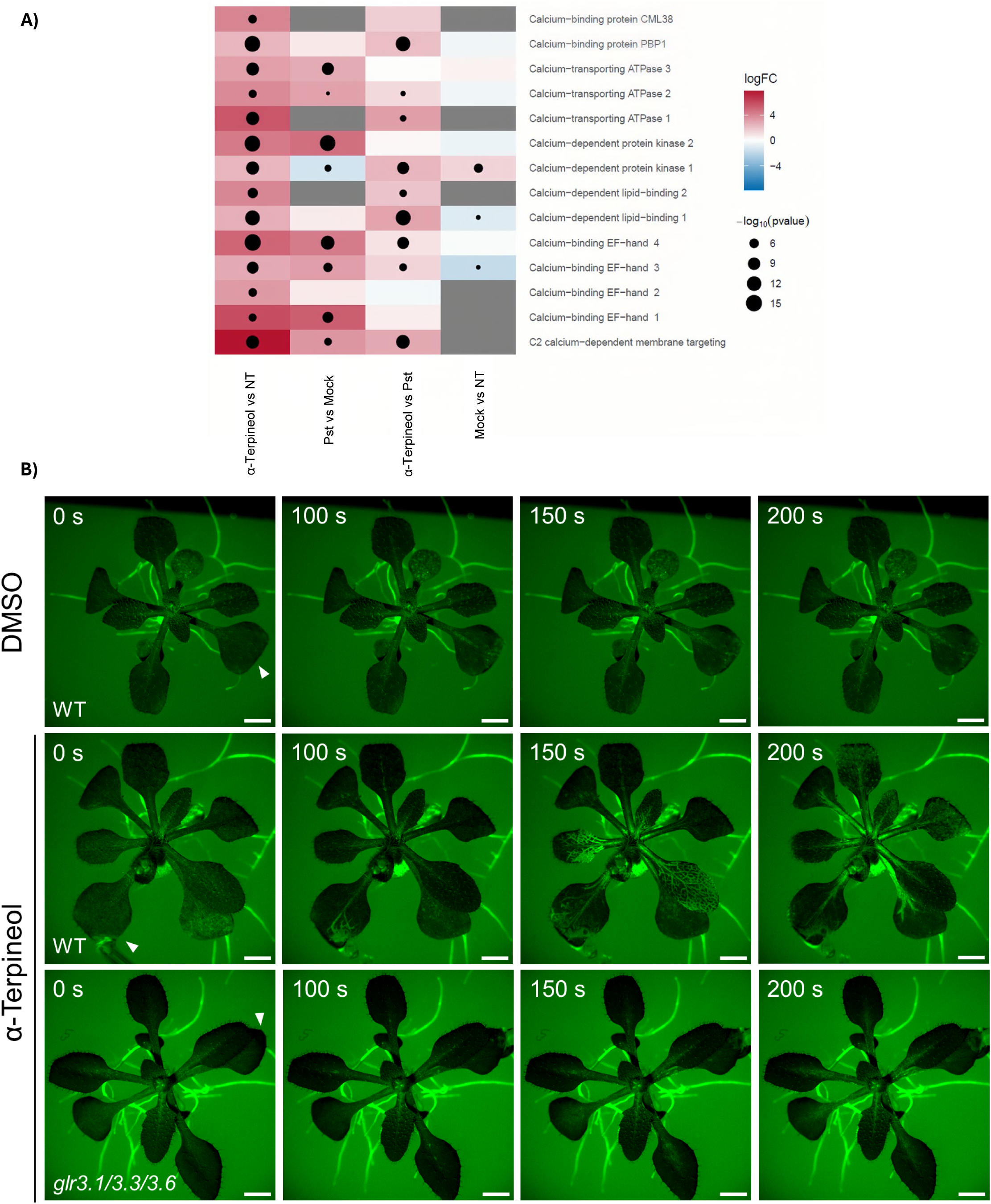
Calcium-related gene expression and calcium imaging following α-terpineol treatment. **(A)** Heatmap representation of differentially expressed genes associated with calcium signalling, including calcium-binding proteins, calcium transporters and calcium-dependent kinases, in response to terpineol treatment compared with *Pseudomonas syringae* infection and mock controls. Colour scale indicates log fold change (logFC), and dot size represents statistical significance (−log *p*-value). **(B)** Representative fluorescence images of 14-day-old *Arabidopsis thaliana* plants constitutively expressing the cytosolic calcium sensor GCaMP3 in the wild-type (WT, *Col-0*) and *glr3.1/3.3/3.6* triple knock-out mutant backgrounds following treatment with DMSO or α-terpineol. White arrowheads indicate the point of compound application. Scale bar: 2 mm.

### α-Terpineol Orchestrates Multilayered Plant Immunity through Hormonal and Metabolic Reprogramming

As shown in Fig. 1C, α-terpineol treatment triggered a significant overrepresentation of GO terms related to “salicylic acid mediated signalling pathway”, “jasmonic acid biosynthetic process”, and “isopentenyl diphosphate biosynthetic process”. The activation of these pathways, classically associated with systemic resistance and specialized metabolism, provided a clear rationale to further dissect the hormonal landscape of the α-terpineol-induced response.

Transcriptomic data revealed the activation of JA and SA biosynthetic and signalling pathways. JA biosynthetic genes (*SlLOX8, SlAOS2*) and key signalling components (*SlMYC2*) were strongly up-regulated (Fig. 4A). Similarly, SA signalling genes (PR proteins; Fig. 4B) and gibberellin signalling genes (Fig. 4C) exhibited pronounced activation. Hormonal quantification confirmed that JA, SA, and gibberellins GA1 and GA4 significantly accumulated following α-terpineol treatment (Fig. 4D-F).

**Figure 4.**
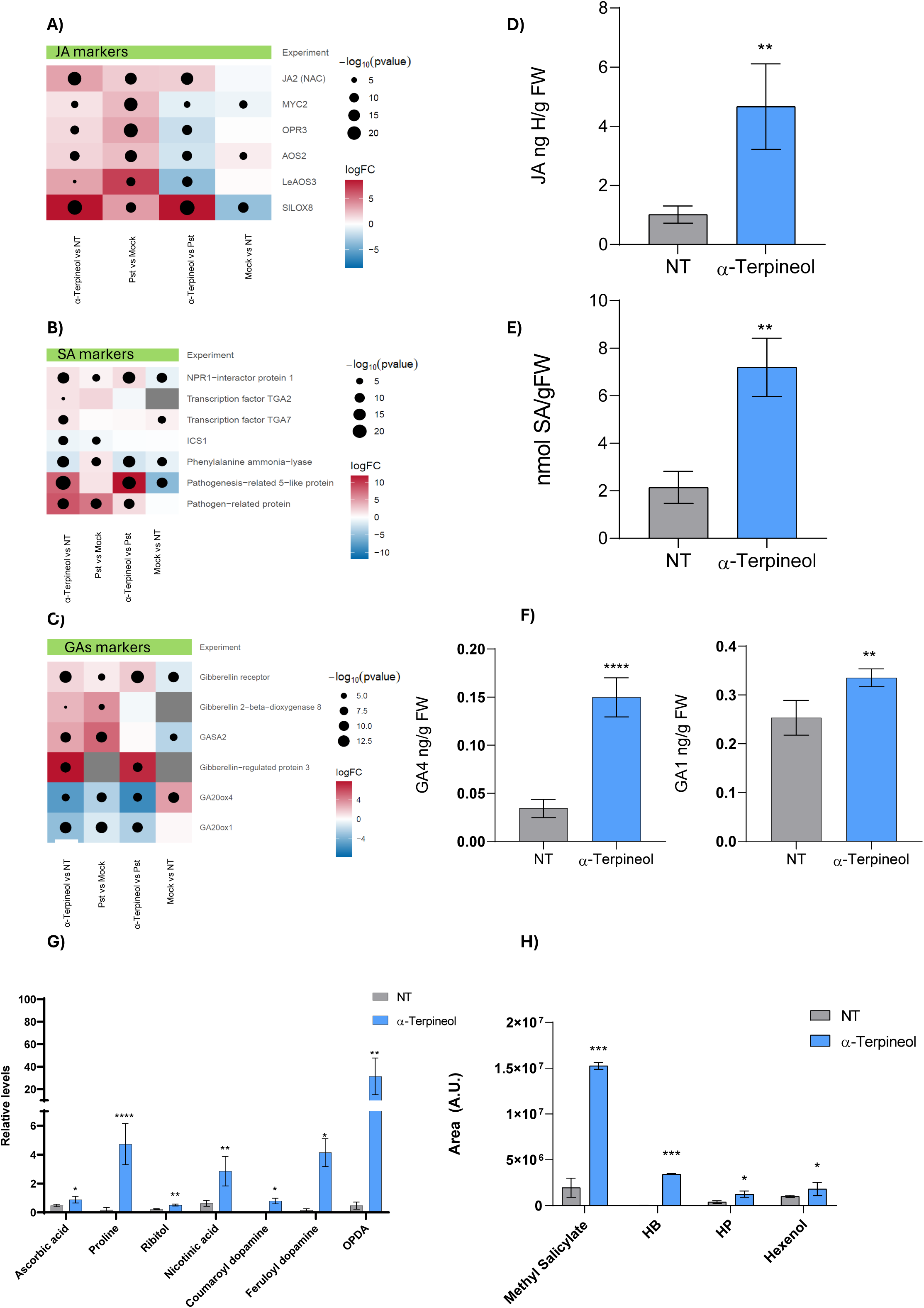
Transcriptomic, hormonal and secondary metabolites analyses in α-terpineol-treated plants. Heatmaps on the left **(A, B** and **C)** show the differential expression of marker genes associated with the jasmonic acid (JA) and salicylic acid (SA) pathways. Colour scales represent log fold change (logFC), and dot size indicates statistical significance (−log *p*-value) for the different experimental comparisons shown below each heatmap. Bar graphs on the right **(D, E** and **F)** display the quantification of phytohormones in non-treated (NT) and α-terpineol-treated plants, including jasmonic acid (JA), salicylic acid (SA), and the bioactive gibberellins GA₁ and GA₄. Hormone levels are expressed as ng g⁻¹ fresh weight (FW) or nmol g⁻¹ FW, as indicated on each axis. Data are shown as mean ± standard error (SE). Statistical significance between treatments is indicated by asterisks (* *p* < 0.05; ** *p* < 0.01; **** *p* < 0.0001). **(G)** Relative levels of secondary metabolites were quantified in non-treated (NT) and in α-terpineol-treated plants. The analysed compounds include phenylpropanoid derivatives (coumaroyl dopamine and feruloyl dopamine), the jasmonic acid precursor OPDA, antioxidants and osmoprotectors (ascorbic acid, proline, and ribitol), as well as redox-related metabolites (nicotinic acid). Metabolite levels are expressed as ratios relative to formononetin. **(H)** GC–MS analysis showing the accumulation of defence-related volatile organic compounds, including methyl salicylate (MeSA), (*Z*)-3-hexenyl butyrate (HB), (*Z*)-3-hexenyl propionate (HP), and hexenol, in leaves of tomato plants treated with α-terpineol compared to non-treated (NT) plants. Data are expressed as peak area in arbitrary units (A.U.).

To further examine whether α-terpineol treatment also reshapes specialized metabolism beyond hormone accumulation, we performed UPLC-MS coupled with an ORBITRAP mass spectrometer. Metabolomic profiling revealed a significant increase in specific metabolites in treated plants compared to controls (Fig. 4G), including ascorbic acid, proline, ribitol, nicotinic acid, and phenylpropanoid derivatives (coumaroyl-dopamine and feruloyl-dopamine), suggesting an enhanced redox and defence-related metabolic activity and reinforcing cell walls to create physical barriers against pathogens (Zeiss et al., 2021). A significant accumulation of 12-oxo-phytodienoic acid (OPDA), a key precursor of JA, was also observed. Analysis of volatile emissions using PCA and hierarchical clustering (Supplementary information, Fig. S5) revealed a broad metabolic reconfiguration, characterized by increased accumulation of green leaf volatiles (GLVs), one of the major classes of defence-related volatiles associated with Pst infection (López-Gresa, 2017; López-Gresa et al., 2018; Payá et al., 2024), together with enhanced emission of methyl salicylate (MeSA) (Fig. 4H). These results support the notion that HMTPs drive coordinated changes in volatile emissions. Taken together, this multilayered reconfiguration suggests that α-terpineol acts as a potent immune-modulating volatile that primes plants for enhanced resistance through the coordinated activation of hormonal signalling and specialized metabolism.

### Jasmonic Acid Signalling Regulates HMTP Biosynthesis During Plant Defence

Once the essential downstream components required for α-terpineol activity were identified, we next sought to determine the upstream signalling pathways regulating their biosynthesis. Since HMTP production is induced upon avirulent Pst infection triggering ETI (López Gresa et al., 2017), we analysed the contribution of defence-associated signalling pathways to HMTP regulation by applying chemical inhibitors *in planta*. Treatments with NAC, EGTA, and DPI significantly reduced HMTP content (Fig. 5A), while no reduction was observed using PD98059 or SHAM (Supplementary information, Fig. S6), indicating that Ca^2+^ and ROS act as key upstream signals.

**Figure 5.**
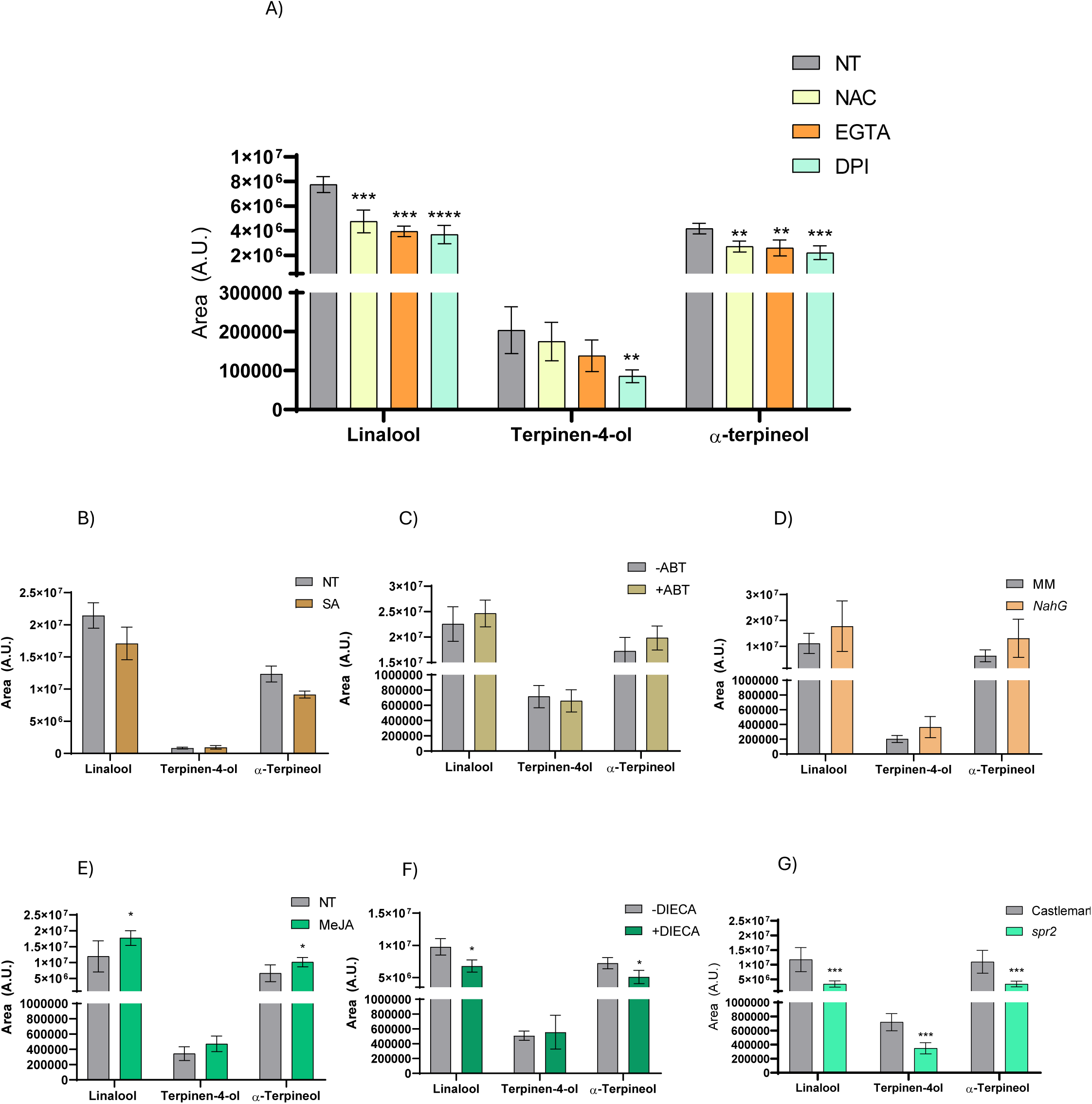
Increase of HMTPs levels in response to salicylic acid, jasmonic acid, and reactive oxygen species treatments. Accumulation of linalool, α-terpineol, and terpine-4-ol was quantified as peak area (arbitrary units, A.U.) under the indicated conditions. **(A)** Prior to infection with *Pseudomonas syringae* pv. *tomato*, *in planta* treatments with chemical inhibitors (NAC, EGTA, or DPI) were performed, with untreated RG tomato plants used as controls (NT). **(B)** HMTP levels in MM tomato plants treated with salicylic acid (SA) compared with non-treated (NT) controls. **(C)** HMTPs in infected RG plants treated with ABT, a SA biosynthesis inhibitor. **(D)** HMTP levels in wild-type MM compared to SA-deficient *NahG* tomato plants. **(E)** HMTP levels in MM tomato plants treated with methyl jasmonate (MeJA) compared with non-treated controls. **(F)** HMTPs in infected RG plants treated DIECA, a JA signalling inhibitor. **(G)** HMTP levels in wild-type Castlemart and JA-deficient *spr2* tomato plants. Data represent mean ± standard error. Statistical differences relative to NT are indicated by asterisks (* *p* < 0.05; ** *p* < 0.01; **** *p* < 0.0001).

Calcium and ROS are well-known mediators of plant defence responses associated with both SA (Herrera-Vásquez et al., 2015; Du et al., 2009) and JA (Hu et al., 2022; Kadam and Barvkar, 2024), two pivotal plant hormones essential for the activation of defence responses. Both hormones also accumulate in contexts where HMTPs are emitted (López Gresa et al., 2017). Exogenous SA treatments did not induce HMTPs accumulation (Fig. 5B) and the application of the SA inhibitor ABT (Koramutla et al.,2022) likewise had no effect (Fig. 5C), and *NahG* tomato plants, which are impaired in SA accumulation, produced HMTPs at levels comparable to wild-type plants (Fig. 5D), demonstrating that SA does not play a relevant role in HMTPs biosynthesis. In contrast, exogenous application of methyl-jasmonate (MeJA) promoted HMTPs biosynthesis (Fig. 5E), whereas the JA inhibitor DIECA (Li et al., 2020) reduced HMTP accumulation (Fig. 5F). Consistently, *spr2* tomato plants, which are defective in JA biosynthesis (Li et al., 2003), showed reduced HMTPs levels (Fig. 5G), identifying JA as the major hormonal regulator of HMTP accumulation (van Schie et al., 2007; Bin et al., 2023). Overall, the data suggest that calcium- and ROS-mediated signalling could converge on JA as the central hormonal regulator of HMTP biosynthesis during plant defence, within the context of ETI.

### HMTPs Establish a Self-Reinforcing Feedback Loop Through Biosynthetic Activation and Chemical Interconversion

In addition to triggering a transcriptional reprograming that resembles Pst infection, α-terpineol promotes extensive metabolic remodelling and induces the propagation of calcium signalling dynamics typical of pathogen-triggered immune responses. Furthermore, α-terpineol requires key defence-associated signalling components, including Ca²⁺, ROS and JA, for both HMTP biosynthesis and downstream signalling. Collectively, these observations suggest that HMTPs may reinforce their own production through a self-amplifying positive feedback mechanism.

To test whether HMTPs promote their own accumulation, plants were treated with α-terpineol or linalool. α-Terpineol treatment induced the synthesis of terpinen-4-ol, whereas linalool application led to increased levels of both terpinen-4-ol and α-terpineol (Fig. 6A).

**Figure 6.**
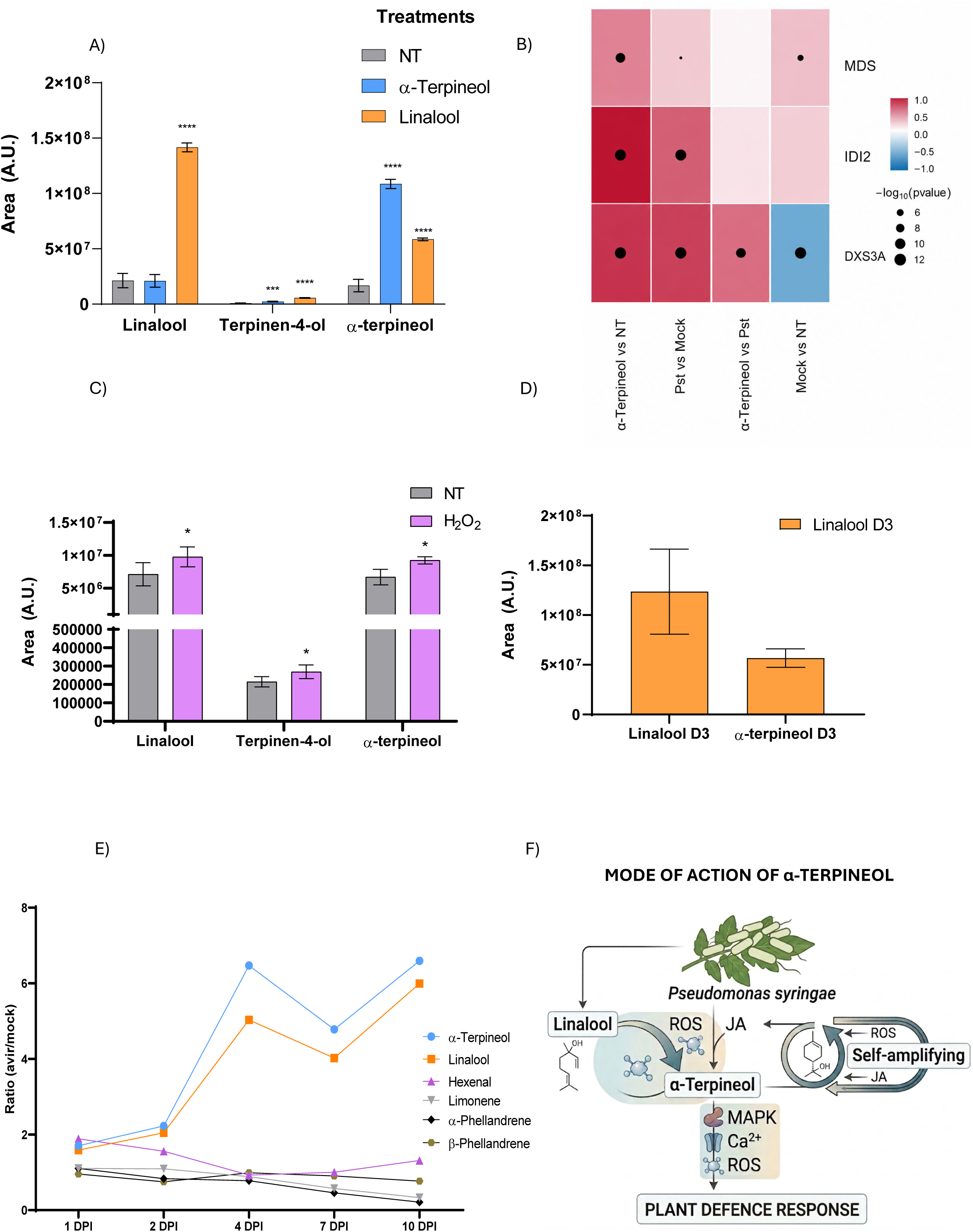
Effects of oxidative conditions on HMTPs accumulation and temporal dynamics of volatile accumulation during avirulent *Pseudomonas syringae* infection. **(A)** Accumulation of linalool, terpinen-4-ol, α-terpineol after α-terpineol or linalool treatments compared with non treated MM tomato plants **(B)** Heatmap representation of differentially expressed genes associated with MEP pathway biosynthesis genes, in response to terpineol treatment compared with *Pseudomonas syringae* infection and mock controls. Colour scale indicates log fold change (logFC), and dot size represents statistical significance (−log p-value). **(C)** Accumulation of linalool, terpinen-4-ol, and α-terpineol in planta after ROS (hydrogen peroxide H₂O₂) treatments compared to non-treated (NT) controls. **(D)** Accumulation of deuterium- labelled monoterpenoids following incubation with linalool-D3, including detection of linalool-D3 and α-terpineol-D3 (RT 30.48). **(E)** Relative accumulation of hydroxylated monoterpenes (α-terpineol and linalool) and non-hydroxylated volatiles (hexenal, limonene, α-phellandrene and β-phellandrene) at 1, 2, 4, 7, and 10 days post-inoculation (DPI). Data are shown as ratios between avirulent- and mock-infected plants. **(F)** The diagram illustrates the molecular mechanism by which the plant responds to *P.syringae* infection. Following pathogen detection (depicted on the leaf at the top), complex signalling pathways are activated. On the left, the synthesis of the hydroxylated monoterpene Linalool (chemical structure shown) and its conversion to α-terpineol is mediated by Reactive Oxygen Species (ROS). α-Terpineol serves as a central signalling node. On the right, a “Self-amplifying” loop involving Jasmonic Acid (JA) and ROS acting as a positive feedback mechanism to sustain the response. Downstream α-terpineol triggers a signalling cascade characterized by the activation MPK, calcium influx, and a secondary wave of ROS production. This coordinated cascade ultimately culminates in the PLANT DEFENCE RESPONSE (bottom box). Arrows indicate activation or flow; chemical structures and molecular icons represent the key species involved.

In contrast, treatment with limonene (non-hydroxylated structure) did not result in significant changes in HMTPs accumulation, indicating that limonene does not undergo interconversion under these conditions (Supplementary information, Fig. S7A). Transcriptomic analysis revealed a strong induction of genes associated with the MEP pathway *MDS*, *IDI2*, and *DXS3a* (Fig. 6B), supporting a positive biosynthetic feedback loop. Furthermore, exogenous hydrogen peroxide (H2O2) treatment in tomato plants significantly increased the levels of α-terpineol, linalool, and terpinen-4-ol (Fig. 6C), indicating that oxidative conditions favour HMTPs production and diversification.

To directly assess whether HMTPs diversification could occur through chemical interconversion *in planta*, tomato plants were treated with deuterium-labelled linalool (linalool-d₃). Remarkably, linalool-d₃ treatment resulted in a significant increase in α-terpineol-d₃, supporting the hypothesis that linalool can be incorporated to the plant and chemically converted into α-terpineol (Fig. 6D).

After establishing that exogenous linalool application specifically promotes α-terpineol accumulation (Fig. 6A) and that exogenous H₂O₂ treatment enhances HMTP levels *in planta* (Fig. 6C), we sought to determine whether oxidative conditions could directly mediate HMTP interconversion. *In vitro* exposure of linalool solutions to H₂O₂ resulted in a substantial decrease in linalool levels together with the appearance of several monoterpenoid-derived peaks (representative ion: *m/z* 93; Supplementary information, Fig. S7B). Among these compounds, α-terpineol (retention time: 30.395 min) was detected exclusively in H₂O₂-treated samples and was completely absent from untreated linalool controls. These results provide evidence that oxidative conditions associated to plant stress may be sufficient to drive the non-enzymatic conversion of linalool into other HMTPs, particularly α-terpineol, supporting a chemical mechanism underlying HMTP diversification during the oxidative burst associated with pathogen defence.

Collectively, these results indicate that HMTP accumulation is sustained through a dual positive feedback mechanism linking transcriptional activation of monoterpenoid biosynthesis with chemical diversification of HMTP-derived compounds. Overall, these findings uncover an additional layer of chemical flexibility shaping volatile-mediated plant immune responses.

### Temporal Dynamics Support Sustained Self-Reinforcing HMTP Accumulation During Plant Defence

Because HMTPs mimic infection-associated responses and promote their own accumulation, we reasoned that HMTPs production could be sustained through self-reinforcing regulation during infection. To determine the extent of this reinforcement, we studied the kinetics of HMTPs accumulation during the avirulent interaction by calculating the ratio between avirulent- and mock-infected plants at different time points post-Pst inoculation (Fig. 6E). HMTPs were differentially emitted, with both α-terpineol and linalool displaying a progressive increase over time, reaching their highest levels at later stages of infection (10 days post-inoculation). In contrast, other volatiles such as hexenal and the monoterpenes limonene and phellandrene showed comparatively minor or declining changes over the same period. This sustained accumulation of HMTPs at late infection stages during avirulent interactions is consistent with a model in which HMTPs production is reinforced over time rather than being transiently induced. This temporal pattern is compatible with defence responses that persist during prolonged plant-pathogen interactions, such as long-lasting systemic acquired resistance, classically associated with avirulent infection. To evaluate how this sustained accumulation is initiated, we also analysed HMTPs dynamics at early time points during avirulent Pst interactions. Our analysis revealed that α-terpineol and linalool levels begin to increase within the first 18 hours after infection and continue to rise steadily up to 48 hours, whereas hexenal and monoterpenes limonene and phellandrene levels remained largely unchanged over this period (Supplementary information, Fig. S8).To better determine whether the observed increase in HMTP levels reflected active production rather than passive retention of exogenously applied volatiles, plants were subjected to individual HMTP treatments and maintained either in sealed boxes or under open-air conditions, allowing volatile dispersion into the environment. As shown in Supplementary information, Fig. S9, comparable levels of α-terpineol, linalool and terpinen-4-ol were detected under both conditions at 24 h and 72 h after treatment. Consistent with the sustained accumulation of HMTPs during ETI, *TPS5* expression remained elevated at early infection stages, whereas MEP pathway genes displayed a transient induction pattern (Supplementary information, Fig. S10), suggesting that long-term HMTP persistence may involve both early biosynthetic activation and subsequent chemical diversification mechanisms. Taken together, our results support a self-reinforcing mechanism shaping HMTPs profiles through transcriptional activation and chemical interconversion during plant immunity (Fig. 6F).

## DISCUSSION

Hydroxylated monoterpenes (HMTPs) have emerged as central players in plant defence mechanisms (Riedlmeier et al., 2017; Wenig et al., 2019; Vlot et al., 2021; Pérez-Pérez et al., 2024), yet their functional integration within immune signalling networks has remained poorly defined. Here, we show that α-terpineol mimics key features of *Pseudomonas syringae* (Pst) infection, while HMTP biosynthesis depends on ROS and jasmonate signalling. Oxidative conditions further promote HMTP diversification, favouring accumulation of the highly bioactive α-terpineol. Together with the ability of HMTPs to promote their own accumulation, these findings support a self-reinforcing immune circuit coupling pathogen mimicry, JA-dependent volatile production and redox-driven chemical diversification during antibacterial defence.

At the molecular level, α-terpineol treatment induces an outstanding transcriptional reprogramming, displaying significant overlap with genes regulated during Pst infection (Fig. 1A), suggesting that HMTPs act as molecular mimics of pathogen-triggered defence responses. Intriguingly, the number of genes regulated by α-terpineol exceeds those regulated during Pst infection, suggesting additional regulatory functions for this compound. GO analysis (Fig. 1C-D) confirms the enrichment of terms related to defence responses and specialized metabolism, consistent with reports linking volatile signalling to metabolic reprogramming (Heil and Silva Bueno, 2007; Scala et al., 2013). This is in agreement with previous works describing defence-associated volatiles as endogenous danger signals capable of activating canonical immune pathways in the absence of pathogen-derived molecules (Payá et al., 2024).

One of the earliest defence responses limiting bacterial entry is stomatal closure, a process tightly regulated by reactive oxygen species (ROS), cytosolic Ca²⁺ fluxes, and mitogen-activated protein kinase (MPK) cascades (Melotto et al., 2006; Kollist et al., 2014). Our results demonstrate that α-terpineol efficiently triggers stomatal closure through signalling pathways involving calcium, NADPH oxidase-dependent MPK activation, and ROS production (Fig. 2). Notably, α-terpineol-induced stomatal closure operates independently of abscisic acid (ABA) (Pérez-Pérez et al., 2024). ABA-independent stomatal immunity has been described in several biotic stress contexts and is frequently mediated by ROS- and Ca²⁺-dependent signalling networks (Montillet et al., 2013; Arnaud and Hwang, 2015; Payá et al. 2024), placing HMTPs within this alternative defensive framework. Importantly, the ability of α-terpineol to propagate cytosolic Ca^2+^ waves across distal tissues (Fig. 3) suggests participation in long-distance defence signalling, consistent with previous studies identifying calcium waves as central mediators of systemic immune communication (Toyota et al., 2018; Wang and Luan, 2024). Together, these observations highlight the dual role of α-terpineol as both a direct immune signal and a potent amplifier of antibacterial defence responses.

The hormonal profiling revealed significant accumulations of jasmonic acid (JA) and salicylic acid (SA) (Fig. 4D, E), demonstrating a broad-spectrum hormonal response. Beyond this, α-terpineol induces extensive remodelling of specialized metabolism (Fig. 4G and 4H), leading to the accumulation of antioxidant and stress-protective metabolites (ascorbic acid, ribitol, proline) and cell wall-reinforcing phenylpropanoids (Campos 2014; Zeiss et al., 2021). In parallel, α-terpineol promotes the accumulation of volatiles associated with systemic acquired resistance (SAR), such as methyl salicylate and green leaf volatiles. The simultaneous induction of these metabolites is particularly striking because it indicates that α-terpineol does not activate an isolated defence branch, but rather orchestrates a multilayered immune state integrating antibacterial, wound-related, and systemic signalling pathways. Pharmacological and genetic approaches demonstrate that both ROS and JA signalling are required for HMTP accumulation during defence (Fig. 5). This interpretation is consistent with previous studies showing JA-dependent induction of terpene synthases and monoterpene emission in tomato trichomes (van Schie et al., 2007) as well as JA-mediated activation of terpene biosynthesis during plant defence responses (Martin et al., 2002; Miller et al., 2005; Bin et al., 2023). In the context of ETI, this observation is particularly relevant, since JA has traditionally been viewed as antagonistic to SA-dependent antibacterial responses but is increasingly recognized as an important component of systemic immunity following pathogen challenge (van Schie et al., 2007). Importantly, the bidirectional relationship between JA signalling and HMTP production establishes a self-reinforcing immune circuit where jasmonates promote HMTP biosynthesis, and HMTPs in turn amplify immune signalling and metabolic reprogramming. These findings support a revised framework in which JA-dependent volatile signalling cooperates with canonical SA pathways through the mediation of HMTPs to coordinate local and systemic antibacterial immunity, where HMTPs act as key mediators.

A particularly novel aspect of our study is the identification of chemical remodelling as a mechanism contributing to HMTPs diversification and bioactivity. Our results indicate that oxidative conditions are sufficient to promote the enzyme-free chemical interconversion of hydroxylated monoterpenes *in vitro*, and deuterium-labelled linalool was converted into deuterated α-terpineol *in planta*. To date, no terpene synthase has been identified in tomato to produce α-terpineol (Zhou and Pichersky, 2020). Although ROS are well-established signal molecules in plant immunity, particularly during ETI establishment (Apel and Hirt, 2004) -a stage in which HMTPs are differentially emitted (López-Gresa et al., 2017), the potential contribution of oxidative stress to reshaping volatile metabolite composition through chemical interconversion has received limited attention. Our findings extend previous observations describing oxidative rearrangements of terpenoids under stress conditions (Attaway et al., 1968; Maffei et al., 2012; Scurria et al., 2020) and reveal an alternate regulatory layer operating alongside canonical biosynthetic pathways to enhance the versatility of the plant response. Notably, oxidative remodelling preferentially favoured the accumulation of α-terpineol, which displays the strongest antibacterial activity among the monoterpenoids tested, thereby linking chemical diversification with functional immune output. Our results indicate that the oxidative conditions accompanying the plant response to pathogens may contribute not only to immune signalling but also to modulation of volatile composition during plant defence. While ROS-driven chemical interconversion appears sufficient to account for α-terpineol formation, additional undescribed enzymatic contributions under specific *in vivo* conditions cannot be ruled out.

Beyond chemical diversification, our data indicate that HMTPs participate in a self-reinforcing immune circuit. Temporal analyses revealed a progressive and persistent accumulation of HMTPs throughout ETI, supporting the existence of a feed-forward mechanism sustaining volatile-mediated defences over time (Fig.s 7A and S8). Together with their ability to activate defence signalling, propagate cytosolic Ca²⁺ waves, and promote their own accumulation, our findings show that HMTPs reinforce immune signalling during infection. Notably, the sustained accumulation of HMTPs resembles long-lasting defence states such as systemic acquired resistance (SAR), which is established following avirulent interactions and maintained over extended periods (Wildermuth et al., 2001; Ding et al., 2020). In this context, HMTPs may contribute to maintain the SAR-associated signalling by sustaining JA-dependent defence programs beyond the initial infection site, which is consistent with recent evidence linking JA signalling to SAR establishment following ETI (Gaikwad et al., 2026).

Collectively, our data support a model in which pathogen perception activates ROS and JA signalling (Wildermuth et al., 2001; Ding et al., 2020), promoting the biosynthesis and accumulation of HMTPs during defence responses (Fig. 6F). Concurrently, the oxidative environment associated with immune activation favours chemical remodelling within HMTP pools, selectively enriching highly bioactive compounds such as α-terpineol. Once accumulated, α-terpineol activates MPK-, ROS- and Ca²⁺-dependent signalling pathways, triggers defence-associated transcriptional, metabolic and volatile reprogramming, and propagates cytosolic Ca²⁺ waves across distal tissues. Simultaneously, HMTPs promote their own accumulation through a JA-dependent feed-forward loop, thereby sustaining volatile production and immune activation over time. In this context, HMTPs may contribute not only to local antibacterial responses but also to the maintenance and amplification of long-lasting systemic immune signalling. Together, this self-reinforcing circuit positions HMTPs as dynamic amplifiers of tomato immunity, linking immune signalling, redox-driven volatile diversification and long-lasting defence activation during infection.

## METHODS

### Plant Material

For this study, various tomato genotypes were utilized: (i) the parental Rio Grande (RG) line carrying the *Pto* resistance gene, kindly provided by Dr. Selena Giménez (Centro Nacional de Biotecnología, Madrid, Spain); (ii) Money Maker (MM) plants as wild-type tomato for volatile treatments; (iii) *spr2* (Li et al., 2003) tomato mutants and its parental Castlemart, kindly provided by Dr. Concha Gómez-Mena (Instituto de Biología Molecular y Celular de Plantas, UPV-CSIC, Valencia, Spain); (iv) *NahG* plants (Brading et al., 2000) and their parental MM (kindly provided by Prof. Jonathan Jones (The Sainsbury Laboratory, Norwich, United Kingdom). Various *Arabidopsis thaliana* genotypes were used: (i) *Arabidopsis thaliana* bearing the ABA-responsive reporter *pMAPKKK18-LUC*+ (kindly provided by Dr. Jorge Lozano, Instituto de Biología Molecular y Celular de Plantas UPV-CSIC, Valencia, Spain), (ii) *rhob* and *mpk* mutants of *Arabidopsis thaliana*, alongside their parental line, provided by Dr. Lucía Jordá (Centro de Biotecnología y Genómica de Plantas, Madrid, Spain); and (iii) GCaMP3 *Arabidopsis thaliana* plants and the triple mutant *glr3.1/3.3./3.6* were provided by Dr. Chysoula Pantazopoulou (Plant-Environment Signalling, Institute of Environment Biology, Utrecht University, Utrecht, The Netherlands). Seeds were surface-sterilized with a 1:1 sodium hypochlorite:distilled water solution, followed by consecutive washes of 5, 10, and 15 minutes to ensure complete removal of the disinfectant. Germinated seeds were sown in 12-cm-diameter pots containing a mixture of vermiculite and peat. Greenhouse conditions were maintained at approximately 50% relative humidity with a 16/8-hour light/dark cycle at 26 °C.

### Chemical treatments

To investigate the effects of various chemical inhibitors on tomato plants, stem feeding treatments were performed as follows: EGTA (10 mM, pH adjusted to 8) (Tang et al., 2019); N-acetylcysteine (NAC) (5 mM, pH adjusted to 8) (Montillet et al., 2013); MEK-1 inhibitor PD98059 (100 µM) (Payá et al., 2024), diphenyleneiodonium (DPI) (100 µM) (Moloi et al., 2006), 3-amino-1,2,4-triazole (ABT) (100 µM) (Fan et al., 2022) and sodium diethyldithiocarbamate (DIECA) at 2 mM. Plants were embedded via stem feeding with the respective inhibitors for 3 hours, followed by placement in methacrylate chambers and exposure to α-terpineol (5 µM) for 24 hours, after which the plants were infected with the pathogen, and bacterial counts were performed to assess pathogen proliferation. For experiments aimed at analysing volatile emissions, plants were incubated with the inhibitors via stem feeding for 3 hours, subsequently infected with the pathogen, and tissue samples were collected.

### Stomatal aperture assays

Leaf discs from 3- to 4-week-old plants were floated on Murashige and Skoog (MS) medium under light for 3 hours to induce stomatal opening as described in Payá et al.,2024. Elicitors such as 1 μM flg22, 10 μM abscisic acid (ABA), or 50 μM α-terpineol were added for an additional 3 hours. Chemical inhibitors (2 mM salicylhydroxamic acid (SHAM), 20 μM diphenyleneiodonium chloride (DPI), 2 mM EGTA, 20 μM PD98059, and 1 mM NAC) were applied 30 minutes prior to the elicitors. Samples were imaged under a Leica DC5000 microscope, and the aperture ratio (stomatal width/length) was quantified using the ImageJ software (https://imagej.net/ij/).

### Bacterial Strain and Inoculation Procedures

The bacterial strain utilized in this research was *Pseudomonas syringae* pv. *tomato* DC3000 (Pst) bearing deletions in the *avrPto* and *avrPtoB* genes, kindly provided by Dr. Selena Giménez (Centro Nacional de Biotecnología, Madrid, Spain) (Lin and Martin, 2005; Ntoukakis et al., 2009). Bacterial cultivation and plant inoculation followed established protocols (López-Gresa et al., 2018). In summary, inoculation was conducted on 4-week-old tomato plants by dipping into a bacterial suspension brought to an optical density of 0.1 at 600 nm, supplemented with 0.05% Silwet L-77. For colony-forming unit (cfu) quantification, three leaf disks (1 cm² each) were excised and homogenized, and serial dilutions of the infected tissue were plated onto King’s B agar plates containing rifampicin. Plates were incubated at 28 °C for 48 hours before counting the cfu.

### RNA Extraction and RT-qPCR

The RNA extraction and conversion to cDNA of tomato leaves were carried out using column kit based on silica membranes (Macherey-Nagel GmbH, Germany) following the manufacturer’s protocol. cDNA from a microgram of RNA was obtained using a PrimeScript RT reagent kit (Perfect Real Time, Takara Bio Inc., Otsu, Shiga, Japan) following its instructions. qPCRs were performed as previously described (Campos et al. 2014). In each 96-well plate, a reaction took place in a final volume of 10 µL. SYBR R Green PCR Master Mix (Applied Biosystems) was used as the fluorescence marker and actin gene as the endogenous reference gene. Primers are listed in Supplementary Table S1. Relative gene expression levels were calculated using the ΔΔCt method and are represented in the figures as fold changes relative to the corresponding reference condition indicated in each experiment.

### RNAseq

For the transcriptomic analysis, three individual 4-week-old Money Maker tomato plants were placed in methacrylate chambers and treated either with terpineol or distilled water. In parallel, another set of plants was inoculated with *Pseudomonas syringae* pv. tomato DC3000 (0.1 O.D. in 10 mM MgCl₂ and 0.05% Silwet L-77) or with a mock solution containing only 10 mM MgCl₂ and 0.05% Silwet L-77. Leaf samples were collected 24 hours after treatment or infection and used for RNA-seq analysis.

RNA sequencing was performed by Novogene using directional mRNA library preparation with poly(A) enrichment. Sequencing was carried out on a NovaSeq 6000 platform (150 bp paired-end reads), generating approximately 60 million reads and 9 Gb of data per sample.

Illumina reads were pre-processed with the Fastp software to remove low-quality bases and adapter sequences using the following parameters: “--cut_front_window_size 1 --cut_front_mean_quality 30 --cut_front --cut_tail_window_size 1 --cut_tail_mean_quality 30 --cut_tail -l 20”. The filtered reads were then aligned to the SL4.0 tomato genome using STAR with default settings.

Raw read counts were obtained with featureCounts (-t “exon”) using the ITAG4.2-merged annotation set (Fernández et al., 2025), and counts were summarized at the gene level. Gene expression values were normalized as fragments per kilobase of transcript per million mapped reads (FPKM).

Differential expression analysis was performed using the LIMMA R package. Genes with an adjusted *p*-value (Benjamini–Hochberg correction) below 0.05 were considered differentially expressed. Functional enrichment analyses were performed on these genes using the PANTHER Classification System (https://pantherdb.org), with gene lists obtained through the gprofiler2 R package.

### Terpineol atlas and Data Availability

A dedicated web application (TomViz tool) was developed within the PlantaeViz platform. This application, which is accessible at https://plantaeviz.tomsbiolab.com/tomviz/terpineol_atlas, provides an interactive interface for examining the DEA results and the normalized gene expression data for each of the replicates generated in the study. It is developed and deployed on Shiny Server to facilitate the exploration of the results.

RNA-seq datasets are available in NCBI SRA database under the identifier PRJNA1458848.

### Salicylic Acid (SA) Analysis

For the analysis of SA, 0.25 g of frozen, homogenized leaf tissue was extracted in methanol containing 25 mM *o*-anisic acid as an internal standard. The mixture was subjected to 10 minutes of sonication and centrifugation, and the supernatant was dried under nitrogen flow. SA quantification followed the protocol described by Vázquez Prol et al. (2021), with calibration curves created using authentic standards.

### Jasmonic Acid (JA) and Gibberellin Analysis

100 mg of finely ground tissue was extracted in 80% methanol containing 1% acetic acid and appropriate internal standards. Samples were mixed by continuous shaking for 1 h at 4 °C, followed by incubation at −20 °C overnight. Extracts were then centrifuged, and the supernatant was collected and evaporated to dryness under vacuum. The resulting residue was resuspended in 1% acetic acid and purified using a reverse-phase solid-phase extraction cartridge (Oasis HLB, 30 mg sorbent; Waters), following the protocol described by Seo et al. (2011).

Final eluates were evaporated to dryness and reconstituted in 5% acetonitrile containing 1% acetic acid. Hormones were separated by ultra-high-performance liquid chromatography (UHPLC) using a reverse-phase Accucore C18 column (2.6 µm particle size, 100 mm length; Thermo Fisher Scientific) with an acetonitrile gradient containing 0.05% acetic acid at a flow rate of 400 µL min⁻¹. For gibberellins (GAs) and jasmonic acid (JA), the gradient ranged from 2% to 55% acetonitrile over 21 min.

Hormone detection was performed using a Q-Exactive Orbitrap mass spectrometer (Thermo Fisher Scientific) operating in targeted Selected Ion Monitoring (tSIM) mode. Instrument settings included a capillary temperature of 300 °C, S-lens RF level of 70, and a resolution of 70,000. Electrospray ionization was conducted with a spray voltage of 3.0 kV, heater temperature of 150 °C, sheath gas flow rate of 40 units, and auxiliary gas flow rate of 10 units. Acidic hormones were analysed in negative ionization mode. Quantification was carried out using embedded calibration curves and processed with Xcalibur 4.0 and TraceFinder 4.1 SP1 software. Deuterium-labelled standards corresponding to each hormone were used as internal standards (OlChemIm Ltd., Olomouc, Czech Republic).

### Analysis of Specialized Metabolites

Tomato leaves from plants treated with α-terpineol (four biological replicates) and untreated control plants (four biological replicates) were freeze-dried, homogenized, and finely ground under liquid nitrogen to ensure sample uniformity. For metabolite extraction, 3 mg of lyophilized tissue was used. Semi-polar metabolites were extracted with 75% methanol containing 0.1% (v/v) formic acid, supplemented with 0.5 µg mL⁻¹ formononetin (Sigma-Aldrich) as internal standard. Extractions were performed at room temperature with continuous agitation for 30 min using an MM 20 mixer mill (Domel) operating at 20 Hz. Samples were centrifuged at 20,000 *g* for 20 min, and 0.5 mL of the supernatant was transferred to 0.2 µm polytetrafluoroethylene (PTFE) filter vials (Whatman) for subsequent LC-MS analysis.

Chromatographic separation was carried out using a Luna C18 column (100x2 mm, 2.5 µm particle size; Phenomenex) maintained at 40 °C. A binary solvent system was used consisting of (A) acetonitrile and (B) water containing 0.1% formic acid as previously described by Wang et al. (2017)

Mass spectrometric analyses were performed on a Q-Exactive hybrid quadrupole-Orbitrap mass spectrometer (Thermo Fisher Scientific, USA) coupled to an Ultimate 3000 HPLC-DAD system. Semi-polar metabolites were analysed using a heated electrospray ionization (HESI) source operated in both positive and negative ionization modes. Data-dependent MS/MS (dd-MS²) fragmentation was applied for metabolite identification. Four replicates were used and the statistical analysis employed was One-way ANOVA comparing the volatiles treatment to the control plants.

Relative quantification was performed by normalizing metabolite peak area to the internal standard. Metabolite identification was based on accurate mass, retention time, and fragmentation spectra, supported by comparisons with authentic standards when available, public spectral libraries, *in-house* databases, and previously published metabolomic data. Accurate mass information was verified using PubChem’s monoisotopic mass database.

Data processing and quantification were performed using Xcalibur Quan (v4.7) software (Thermo Fisher Scientific).

### Volatile Organic Compound (VOC) Analysis

For VOC analysis, 1 mL of 6 M CaCl2 and 100 µL of 750 mM EDTA (pH 7.5) were combined with 100 mg of ground tomato leaves in a 10-mL glass vial. The vials were sealed tightly and subjected to 5 minutes of sonication. Volatile compounds were extracted using headspace solid-phase microextraction (HS-SPME) as described by López-Gresa et al., 2017. Chromatograms and mass spectra were analysed with Enhanced ChemStation software (Agilent), which includes an integrated database for matching ion patterns and retention times to reference compounds. Monoterpene quantification was carried out using an Agilent-developed method that employed the most prominent ions and retention times for peak area calculations. The quantifier ions (*m/z*) and retention times used for VOC measurement were as follows: (i) α-terpineol: *m/z* 93/121 at 30.6 min; (ii) linalool: *m/z* 55/43 at 27.02 min; and (iii) terpinen-4-ol: *m/z* 93/111 at 30.2 min.

For the time-course analysis of volatile production, five plants per condition and time point were used. At day 0, half of the plants were inoculated with the avirulent *Pseudomonas syringae* pv. *tomato* (Pst) strain, while the remaining plants were mock-treated. Samples were collected at 1, 2, 4, 7, and 10 days post-inoculation (dpi). At each time point, five infected plants and five mock-treated plants were harvested. For consistency, the third and fourth leaves, which had been marked at the beginning of the experiment, were collected. Data are presented as the ratio of volatile accumulation in avirulent-infected plants relative to mock-treated controls.

### GCaMP3 Fluorescence Visualization

Arabidopsis plants expressing the genetically encoded Ca^2+^ sensor GCaMP3 were surface sterilized with bleach and stratified at 4 °C for 2 days. Plants were then grown on 1X Murashige and Skoog (MS) agar plates in a controlled environment at 22 °C under a 16 h/8 h light/dark cycle. Imaging was performed using a Leica M205 FA stereo microscope equipped with an sCMOS camera (Leica DFC9000 GT) and a LEICA P eGFP Ex470/40 filter. Images were acquired with a Plan Apo 1X objective using an exposure time of 0.75 s and 3×3 camera binning.

To elicit calcium responses, 1 μl of α-terpineol was applied to the tip of the L1 leaf from 14-day-old Arabidopsis plants. DMSO was used as a control treatment. Before imaging, plants were exposed to blue light for 2 minutes to allow acclimation. Images were analysed using the Fiji package (https://imagej.net/software/fiji/).

### Peroxidase and Superoxide Dismutase Assay

Tomato leaves from plants treated with α-terpineol (four biological replicates) and untreated control plants (four biological replicates) were harvested, immediately frozen in liquid nitrogen, and stored until use. Frozen leaf tissue was finely ground and homogenized in extraction buffer containing 0.1 M Tris (pH 7.0), 0.1% ascorbic acid, 0.1% L-cysteine, 0.5 M sucrose, and 10 mg mL⁻¹ polyvinylpyrrolidone (PVP). Homogenates were centrifuged at 4°C for 15 min, and the resulting supernatants were collected for further analys. Total soluble protein content was determined using the Bradford assay as described in Salazar-Sarasua et al 2022.

Superoxide dismutase (SOD) activity was assayed by adding 25 µg of protein from the crude extract to 1 mL of SOD reaction buffer containing 50 mM phosphate-buffered saline (PBS, pH 7.6), 0.01 mM EDTA, 50 mM sodium carbonate, 12 mM L-methionine, 10 µM riboflavin, and 50 µM nitroblue tetraazolium (NBT). Samples were incubated at room temperature (22-25 °C) under continuous light for 10 min. Absorbance was recorded at 550 nm, using reaction buffer without extract as a blank. SOD activity was expressed as the amount of enzyme required to inhibit 50% of NBT photoreduction.

Peroxidase (PRX) activity was measured by adding 2-75 µg of protein to 1 mL of PRX assay buffer containing 0.85 mM hydrogen peroxide in HEPES buffer (pH 7.0), 0.125 M 4-aminoantipyrine, and 8.1 mg mL⁻¹ phenol. The increase in absorbance was monitored at 510 nm for 2 min. PRX activity was calculated using a calibration curve generated with known concentrations of horseradish peroxidase.

### Chemical Screening for ABA-Responsive Compounds

Seeds from transgenic *Arabidopsis thaliana* (*pMAPKKK18::LUC+*), kindly provided by Dr. Jorge Lozano (Instituto de Biología Molecular y Celular de Plantas, UPV-CSIC, Valencia, Spain) were sterilized and plated in 96-well plates containing half-strength MS medium. Seven-day-old seedlings were treated with DMSO as negative control, ABA as positive control and α-terpineol at a final concentration of 25 μM along with 100 μM D-luciferin. Luminescence was imaged after 6 hours using a LAS-3000 imager as described in García-Maquilón et al. (2021).

## Acknowledgements

We would like to thank the IBMCP Metabolomics Platform (Valencia, Spain), especially Teresa Caballero, for her excellent technical support in VOC quantification. We also thank Dr. Esther Carrera and Jorge Baños for the hormone quantification carried out at the Plant Hormone Quantification Service, IBMCP, Valencia, Spain. We would like to thank María del Rosario González Bermúdez for her support in the ABA luciferase reporter assay. Finally, we thank Dr. Rafael Vázquez Manrique for his assistance in the use of the Leica M205 FA stereo microscope (Generalitat Valenciana equipment) located at the Hospital La Fe in Valencia, Spain.

## Contributions

J.P.-P. designed and executed the research project, performed most of the experimental work, and contributed to wrote the manuscript. P.B.-G. and J.P.-P. performed the GCaMP3 calcium imaging experiments. A.S. and J.T.M. processed raw RNA-seq data and performed differential gene expression. M.S., J.P-P and G.D. carried out metabolite analyses. F.V.-S. provided experimental support and contributed to data acquisition. M.S. and I.R. provided scientific support, contributed to data interpretation, and assisted in manuscript writing. J.P.-P, M.P.L.-G. and P.L. wrote the manuscript. M.P.L.-G. and P.L. conceived the study and supervised the project. All authors discussed the results, revised the manuscript, and approved the final version.

## REFERENCES

Ali, S., Tyagi, A., and Mir, Z.A. (2024). Plant immunity: At the crossroads of pathogen perception and defence response. Plants 13, 1434. 10.3390/plants13111434.

Apel, K., and Hirt, H. (2004). Reactive oxygen species: Metabolism, oxidative stress, and signal transduction. Annual Review of Plant Biology, 55, 373–399. 10.1146/annurev.arplant.55.031903.141701

Arimura, G., Ozawa, R., Kugimiya, S., Takabayashi, J., and Bohlmann, J. (2004). Herbivore-induced defence response in a model legume: Two-spotted spider mites induce emission of (E)-β-ocimene and transcript accumulation of (E)-β-ocimene synthase in *Lotus japonicus*. Plant Physiology 135, 1976–1983. 10.1104/pp.104.042929.

Arnaud, D., and Hwang, I. (2015). A sophisticated network of signalling pathways regulates stomatal defences to bacterial pathogens. Molecular Plant 8, 566–581. 10.1016/j.molp.2014.10.012.

Attaway, J. A. and Buslig, B. S. (1968). Conversion of linalool to alpha-terpineol in citrus. Biochimica et Biophysica Acta, 164(3), 609–610. 10.1016/0005-2760(68)90194-x

Bellés, J.M., Garro, R., Pallás, V., Fayos, J., Rodrigo, I., and Conejero, V. (2006). Accumulation of gentisic acid as associated with systemic infections but not with the hypersensitive response in plant-pathogen interactions. Planta 223, 500–511. 10.1007/s00425-005-0109-8.

Bin, M., Peng, X., Yi, G., and Zhang, X. (2023). CsTPS21 encodes a jasmonate-responsive monoterpene synthase producing β-ocimene in citrus against Asian citrus psyllid. Plant Physiology and Biochemistry 201, 107887. 10.1016/j.plaphy.2023.107887.

Brading, P. A., Hammond-Kosack, K. E., Parr, A. and Jones, J. D. G. (2000). Salicylic acid is not required for Cf-2- and Cf-9-dependent resistance of tomato to *Cladosporium fulvum*. The Plant Journal, 23(3), 305–318.

Brilli, F., Loreto, F., and Baccelli, I. (2019). Exploiting plant volatile organic compounds (VOCs) in agriculture to improve sustainable defence strategies and productivity of crops. Frontiers in Plant Science 10, 264. 10.3389/fpls.2019.00264.

Campos, L., Granell, P., Tarraga, S., López-Gresa, M.P., Conejero, V., Bellés, J.M., Rodrigo, I., and Lisón, P. (2014). Salicylic acid and gentisic acid induce RNA silencing-related genes and plant resistance to RNA pathogens. Plant Physiology and Biochemistry 77, 35–43. 10.1016/j.plaphy.2014.01.016.

Degenhardt, J., Köllner, T.G., and Gershenzon, J. (2009). Monoterpene and sesquiterpene synthases and the origin of terpene skeletal diversity in plants. Phytochemistry 70, 1621–1637. 10.1016/j.phytochem.2009.07.030.

Ding, P., and Ding, Y. (2020). Stories of salicylic acid: A plant defence hormone. Trends in Plant Science 25, 549–565. 10.1016/j.tplants.2020.01.004.

Du, L., Ali, G.S., Simons, K.A., Hou, J., Yang, T., Reddy, A.S.N., and Poovaiah, B.W. (2009). Ca2+/calmodulin regulates salicylic-acid-mediated plant immunity. Nature 457, 1154–1158. 10.1038/nature07612.

Dudareva, N., Klempien, A., Muhlemann, J.K., and Kaplan, I. (2013). Biosynthesis, function and metabolic engineering of plant volatile organic compounds. New Phytologist 198, 16–32. 10.1111/nph.12145.

Evans, K.V., Ransom, E., Nayakoti, S., Wilding, B., Mohd Salleh, F., Gržina, I., Erber, L., Tse, C., Hill, C., Polanski, K., Holland, A., Bukhat, S., Herbert, R.J., de Graaf, B.H.J., Denby, K., Buchanan-Wollaston, V., and Rogers, H.J. (2024). Expression of the Arabidopsis redox-related LEA protein, SAG21 is regulated by ERF, NAC and WRKY transcription factors. Scientific Reports 14, 7756. 10.1038/s41598-024-58161-0.

Fan, S., Wu, H., Gong, H. and Guo, J. (2022). The salicylic acid mediates selenium-induced tolerance to drought stress in tomato plants. Scientia Horticulturae, 300, 111092. 10.1016/j.scienta.2022.111092

Fernández, J. D., Navarro-Payá, D., Santiago, A., Cerda, A., Canan, J., Contreras-Riquelme, S., Moyano, T. C., Landaeta-Sepúlveda, D., Melet, L., Canales, J., Johnson, N. R., Álvarez, J. M., Matus, J. T. and Vidal, E. A. (2025). Organ-level gene-regulatory networks inferred from transcriptomic data reveal context-specific regulation and highlight novel regulators of ripening and ABA-mediated responses in tomato. Plant Communication, 6(11), 101499. 10.1016/j.xplc.2025.101499

Gaikwad, K., Rambla, J.L., Blanca, J., Granell, A., and Trupkin, S.A. (2026). Rapid local and systemic jasmonate signalling is essential for systemic acquired resistance following effector-triggered immunity. Nature Plants 12, 34–45.

García-Maquilón, I., Rodríguez, P.L., Vaidya, A.S., and Lozano-Juste, J. (2021). A luciferase reporter assay to identify chemical activators of ABA signalling. In Plant Chemical Genomics, G.R. Hicks and C. Zhang, eds. (Springer), Vol. 2213. 10.1007/978-1-0716-0954-5_10.

Gomez, S.K., Cox, M.M., Bede, J.C., Inoue, K., Alborn, H.T., Tumlinson, J.H., and Korth, K.L. (2005). Lepidopteran herbivory and oral factors induce transcripts encoding novel terpene synthases in *Medicago truncatula*. Archives of Insect Biochemistry and Physiology 58, 114–127. 10.1002/arch.20037.

González-Guzmán, M., Rodríguez, L., Lorenzo-Orts, L., Pons, C., Sarrión-Perdigones, A., Fernández, M.A., Peirats-Llobet, M., Forment, J., Moreno-Alvero, M., Cutler, S.R., Albert, A., Granell, A., and Rodríguez, P.L. (2014). Tomato PYR/PYL/RCAR abscisic acid receptors show high expression in root, differential sensitivity to the abscisic acid agonist quinabactin, and the capability to enhance plant drought resistance. Journal of Experimental Botany 65, 4451–4464. 10.1093/jxb/eru219.

Heil, M., and Silva Bueno, J.C. (2007). Within-plant signalling by volatiles leads to induction and priming of an indirect plant defence in nature. Proceedings of the National Academy of Sciences of the United States of America 104, 5467–5472. 10.1073/pnas.0610266104.

Herrera-Vásquez, A., Salinas, P., and Holuigue, L. (2015). Salicylic acid and reactive oxygen species interplay in the transcriptional control of defence genes expression. Frontiers in Plant Science 6, 171. 10.3389/fpls.2015.00171.

Howe, G.A., and Jander, G. (2008). Plant immunity to insect herbivores. Annual Review of Plant Biology 59, 41–66. 10.1146/annurev.arplant.59.032607.092825.

Hu, C., Wu, S., Li, J., Dong, H., Zhu, C., Sun, T., Hu, Z., Foyer, C.H., and Yu, J. (2022). Herbivore-induced Ca2+ signals trigger a jasmonate burst by activating ERF16-mediated expression in tomato. New Phytologist 236, 1796–1808. 10.1111/nph.18455.

Kadam, S.B., and Barvkar, V.T. (2024). COI1 dependent jasmonic acid signalling positively modulates ROS scavenging system in transgenic hairy root culture of tomato. Plant Physiology and Biochemistry 206, 108229. 10.1016/j.plaphy.2023.108229.

Karpinska, B., Razak, N., Shaw, D.S., Plumb, W., Van De Slijke, E., Stephens, J., De Jaeger, G., Murcha, M.W., and Foyer, C.H. (2022). Late embryogenesis abundant (LEA)5 regulates translation in mitochondria and chloroplasts to enhance growth and stress tolerance. Frontiers in Plant Science 13, 875799. 10.3389/fpls.2022.875799.

Kollist, H., Nuhkat, M., and Roelfsema, M.R.G. (2014). Closing gaps: Linking elements that control stomatal movement. New Phytologist 203, 44–62. 10.1111/nph.12832.

Koramutla, M.K., Tuan, P.A., and Ayele, B.T. (2022). Salicylic acid enhances adventitious root and aerenchyma formation in wheat under waterlogged conditions. International Journal of Molecular Sciences 23, 1243. 10.3390/ijms23031243.

Li, C., Liu, G., Xu, C., Lee, G.I., Bauer, P., Ling, H.-Q., Ganal, M.W., and Howe, G.A. (2003). The tomato suppressor of prosystemin-mediated responses2 gene encodes a fatty acid desaturase required for the biosynthesis of jasmonic acid and the production of a systemic wound signal for defence gene expression. The Plant Cell 15, 1646–1661. 10.1105/tpc.012237.

Li, M., Yu, G., Cao, C., and Liu, P. (2021). Metabolism, signalling, and transport of jasmonates. Plant Communications 2, 100231. 10.1016/j.xplc.2021.100231.

Li, Y., Qiu, L., Zhang, Q., Zhuansun, X., Li, H., Chen, X., Krugman, T., Sun, Q., and Xie, C. (2020). Exogenous sodium diethyldithiocarbamate, a jasmonic acid biosynthesis inhibitor, induced resistance to powdery mildew in wheat. Plant Direct 4, e00212. 10.1002/pld3.212.

Lin, N. C. and Martin, G. B. (2005). An avrPto/avrPtoB mutant of *Pseudomonas syringae* pv. tomato DC3000 does not elicit Pto-mediated resistance and is less virulent on tomato. Molecular Plant-Microbe Interactions, 18(1), 43–51. 10.1094/MPMI-18-0043

López-Gresa, M.P., Lisón, P., Campos, L., Rodrigo, I., Rambla, J.L., Granell, A., Conejero, V., and Bellés, J.M. (2017). A non-targeted metabolomics approach unravels the VOCs associated with the tomato immune response against *Pseudomonas syringae*. Frontiers in Plant Science 8, 1188. 10.3389/fpls.2017.01188.

López-Gresa, M.P., Payá, C., Ozáez, M., Rodrigo, I., Conejero, V., Klee, H., Bellés, J.M., and Lisón, P. (2018). A new role for green leaf volatile esters in tomato stomatal defense against *Pseudomonas syringae* pv. *tomato*. Frontiers in Plant Science 9, 1855. 10.3389/fpls.2018.01855.

Maffei, M. E., Gertsch, J. and Appendino, G. (2012). Plant volatiles: Production, function and pharmacology. Natural Product Reports, 29(11), 1359–1380. 10.1039/c2np20024g

Martin, D., Tholl, D., Gershenzon, J., and Bohlmann, J. (2002). Methyl jasmonate induces traumatic resin ducts, terpenoid resin biosynthesis, and terpenoid accumulation in developing xylem of Norway spruce stems. Plant Physiology 129, 1003–1018. 10.1104/pp.011001.

Melotto, M., Underwood, W., Koczan, J., Nomura, K., and He, S.Y. (2006). Plant stomata function in innate immunity against bacterial invasion. Cell 126, 969–980. 10.1016/j.cell.2006.06.054.

Miller, B., Madilao, L.L., Ralph, S., and Bohlmann, J. (2005). Insect-induced conifer defence: White pine weevil and methyl jasmonate induce traumatic resinosis and volatile emissions in Sitka spruce. Plant Physiology 137, 369–382. 10.1104/pp.104.050187.

Moloi, M. J. and van der Westhuizen, A. J. (2006). The reactive oxygen species are involved in resistance responses of wheat to the Russian wheat aphid. Journal of Plant Physiology, 163(11), 1118–1125. 10.1016/j.jplph.2005.07.014

Montillet, J.-L., Leonhardt, N., Mondy, S., Tranchimand, S., Rumeau, D., Boudsocq, M., Garcia, A.V., Douki, T., Bigeard, J., Laurière, C., Chevalier, A., Castresana, C., Hirt, H., and Vanacker, H. (2013). An abscisic acid-independent oxylipin pathway controls stomatal closure and immune defence in *Arabidopsis*. PLOS Biology 11, e1001513. 10.1371/journal.pbio.1001513.

Nguyen, C.T., Kurenda, A., Stolz, S., Chételat, A., and Farmer, E.E. (2018). Identification of cell populations necessary for leaf-to-leaf electrical signaling in a wounded plant. Proceedings of the National Academy of Sciences of the United States of America 115, 10178–10183. 10.1073/pnas.1807049115.

Ntoukakis, V., Mucyn, T. S., Gimenez-Ibanez, S., Chapman, H. C., Gutierrez, J. R., Balmuth, A. L., Jones, A. M. and Rathjen, J. P. (2009). Host inhibition of a bacterial virulence effector triggers immunity to infection. Science, 324(5928), 784–787. 10.1126/science.1169430

Pantazopoulou, C.K., Buti, S., Nguyen, C.T., Oskam, L., Weits, D.E., Farmer, E.E., Kajala, K., and Pierik, R. (2023). Mechanodetection of neighbor plants elicits adaptive leaf movements through calcium dynamics. Nature Communications 14, 5827. 10.1038/s41467-023-41530-0.

Payá, C., Belda-Palazón, B., Vera-Sirera, F., Pérez-Pérez, J., Jordá, L., Rodrigo, I., Bellés, J.M., López-Gresa, M.P., and Lisón, P. (2024). Signalling mechanisms and agricultural applications of (Z)-3-hexenyl butyrate-mediated stomatal closure. Horticulture Research 11, uhad248. 10.1093/hr/uhad248.

Pérez-Pérez, J., Minguillón, S., Kabbas-Piñango, E., Payá, C., Campos, L., Rodríguez-Concepción, M., Espinosa-Ruiz, A., Rodrigo, I., Bellés, J.M., López-Gresa, M.P., and Lisón, P. (2024). Metabolic crosstalk between hydroxylated monoterpenes and salicylic acid in tomato defence response against bacteria. Plant Physiology 195, 2323–2338. 10.1093/plphys/kiae148.

Pichersky, E., and Raguso, R.A. (2018). Why do plants produce so many terpenoid compounds? New Phytologist 220, 692–702. 10.1111/nph.14178.

Riedlmeier, M., Ghirardo, A., Wenig, M., Knappe, C., Koch, K., Georgii, E., Dey, S., Parker, J.E., Schnitzler, J.-P., and Vlot, A.C. (2017). Monoterpenes support systemic acquired resistance within and between plants. The Plant Cell 29, 1440–1459. 10.1105/tpc.16.00898.

Ruan, J., Zhou, Y., Zhou, M., Yan, J., Khurshid, M., Weng, W., Cheng, J., and Zhang, K. (2019). Jasmonic acid signalling pathway in plants. International Journal of Molecular Sciences 20, 2479. 10.3390/ijms20102479.

Salazar-Sarasua, B., López-Martín, M. J., Roque, E., Hamza, R., Cañas, L. A., Beltrán, J. P. and Gómez-Mena, C. (2022). The tapetal tissue is essential for the maintenance of redox homeostasis during microgametogenesis in tomato. The Plant Journal, 112(5), 1281–1297. 10.1111/tpj.16014

Scala, A., Allmann, S., Mirabella, R., Haring, M.A., and Schuurink, R.C. (2013). Green leaf volatiles: A plant’s multifunctional weapon against herbivores and pathogens. International Journal of Molecular Sciences 14, 17781–17811. 10.3390/ijms140917781.

Scurria, A., Sciortino, M., Presentato, A., Lino, C., Piacenza, E., Albanese, L., Zabini, F., Meneguzzo, F., Nuzzo, D., Pagliaro, M., Chillura Martino, D. F., Alduina, R., Avellone, G. and Ciriminna, R. (2020). Volatile Compounds of Lemon and Grapefruit IntegroPectin. Preprints, 202012.0034.v1. 10.20944/preprints202012.0034.v1

Seo, M., Jikumaru, Y. and Kamiya, Y. (2011). Profiling of hormones and related metabolites in seed dormancy and germination studies. Methods in Molecular Biology, 773, 99–111. 10.1007/978-1-61779-231-1_7

Tang, Y., Zhao, D., Meng, J. and, et al. (2019). EGTA reduces the inflorescence stem mechanical strength of herbaceous peony. Horticulture Research, 6, 36. 10.1038/s41438-019-0117-7

Toyota, M., Spencer, D., Sawai-Toyota, S., Jiaqi, W., Zhang, T., Koo, A.J., Howe, G.A., and Gilroy, S. (2018). Glutamate triggers long-distance, calcium-based plant defense signaling. Science 361, 1112–1115. 10.1126/science.aat7744.

Van Schie, C.C.N., Haring, M.A., and Schuurink, R.C. (2007). Tomato linalool synthase is induced in trichomes by jasmonic acid. Plant Molecular Biology 64, 251–263. 10.1007/s11103-007-9149-8.

Vázquez Prol, F., Márquez-Molins, J., Rodrigo, I., López-Gresa, M. P., Bellés, J. M., Gómez, G., Pallás, V. and Lisón, P. (2021). Symptom severity, infection progression and plant responses in *Solanum* plants caused by three pospiviroids vary with the inoculation procedure. International Journal of Molecular Sciences, 22(12), 6189. 10.3390/ijms22126189

Vlot, A.C., Sales, J.H., Lenk, M., Bauer, K., Brambilla, A., Sommer, A., Chen, Y., Wenig, M., and Nayem, S. (2021). Systemic propagation of immunity in plants. New Phytologist 229, 1234–1250. 10.1111/nph.16953.

Wang, C., and Luan, S. (2024). Calcium homeostasis and signaling in plant immunity. Current Opinion in Plant Biology 77, 102485. 10.1016/j.pbi.2023.102485.

Wang, C.-C., Sulli, M. and Fu, D.-Q. (2017). The role of phytochromes in regulating biosynthesis of sterol glycoalkaloid in eggplant leaves. PLOS ONE, 12(12), e0189481. 10.1371/journal.pone.0189481

Wenig, M., Ghirardo, A., Sales, J.H., Pabst, E.S., Breitenbach, H.H., Antritter, F., Weber, B., Lange, B., Lenk, M., Cameron, R.K., Schnitzler, J.-P., and Vlot, A.C. (2019). Systemic acquired resistance networks amplify airborne defence cues. Nature Communications 10, 3813. 10.1038/s41467-019-11798-2.

Wildermuth, M.C., Dewdney, J., Wu, G., and Ausubel, F.M. (2001). Isochorismate synthase is required to synthesize salicylic acid for plant defence. Nature 414, 562–565. 10.1038/35107108.

Xiao, Y., Savchenko, T., Baidoo, E.E.K., Chehab, W.E., Hayden, D.M., Tolstikov, V., Corwin, J.A., Kliebenstein, D.J., Keasling, J.D., and Dehesh, K. (2012). Retrograde signalling by the plastidial metabolite MEcPP regulates expression of nuclear stress-response genes. Cell 149, 1525–1535. 10.1016/j.cell.2012.04.038.

Zeiss, D.R., Piater, L.A., and Dubery, I.A. (2021). Hydroxycinnamate amides: Intriguing conjugates of plant protective metabolites. Trends in Plant Science 26, 184–195. 10.1016/j.tplants.2020.09.011.

Zhou, F., and Pichersky, E. (2020). The complete functional characterisation of the terpene synthase family in tomato. New Phytologist 226, 1341–1360. 10.1111/nph.16431.

