## Supporting Material for "Hydroxylated monoterpenes mimic bacterial attack to trigger jasmonate-dependent self-amplifying immunity in tomato"

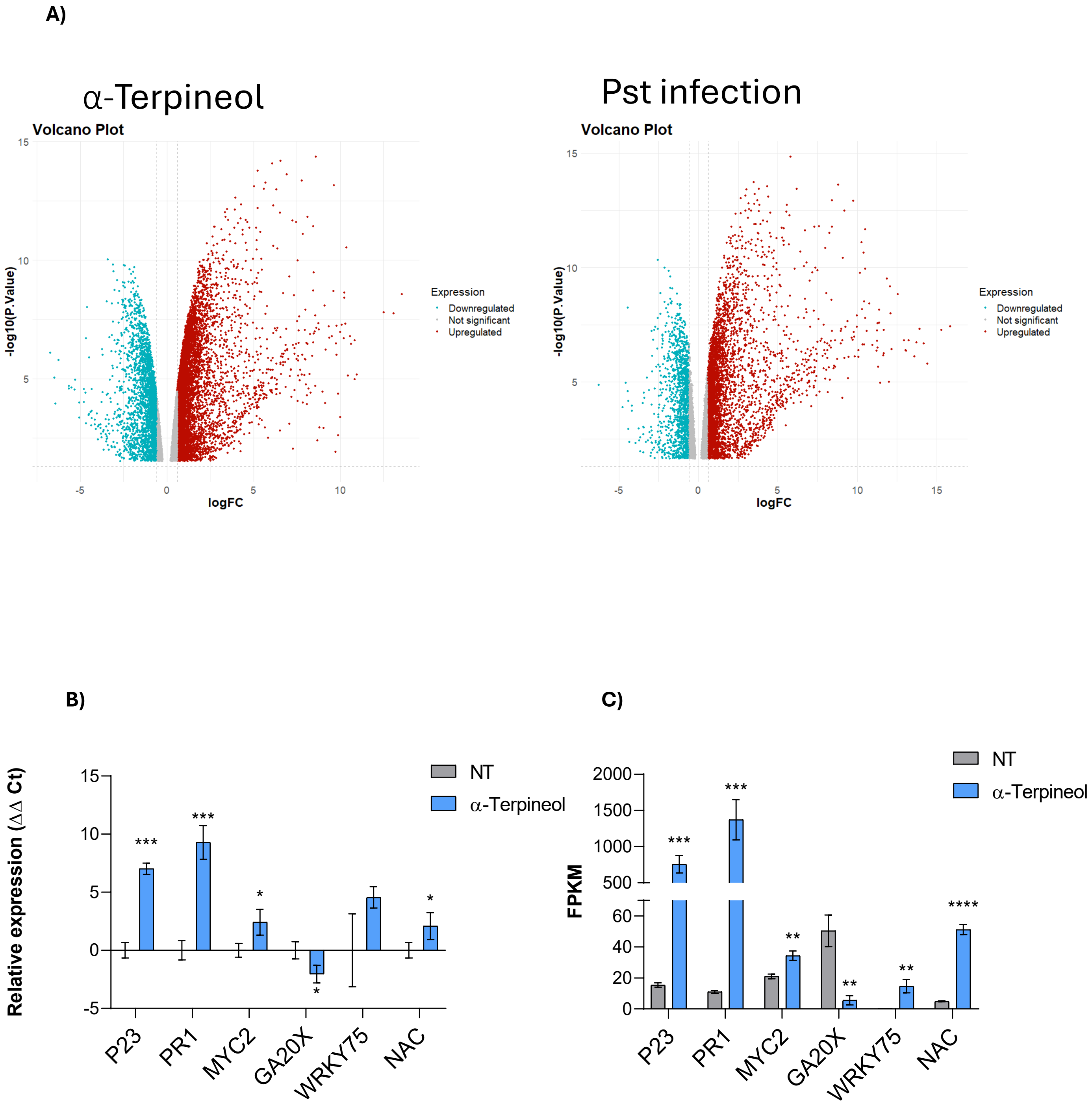


**Figure S1. Transcriptomic analyses of plant responses to α-terpineol treatment and *Pseudomonas syringae* pv. *tomato* (*Pst*) infection.** **(A)** Volcano plots displaying the distribution of differentially expressed genes in response to α-terpineol treatment (left) and Pst infection (right). Genes are plotted according to $\log_2$ fold change and $-\log_{10}$ adjusted p-value, with upregulated, downregulated, and non-significant genes indicated by different colours. Expression levels of selected immune marker genes (*P23, PR1, MYC2, GA2OX, WRKY75,* and *NAC*) in non-treated (NT) and α-terpineol-treated plants were measured by **(B)** RT-qPCR (ΔΔCt) and **(C)** RNA-seq (FPKM) analysis. Data are presented as mean ± standard error (* $p < 0.05$; ** $p < 0.01$, *** $p < 0.001$, **** $p < 0.0001$ ).

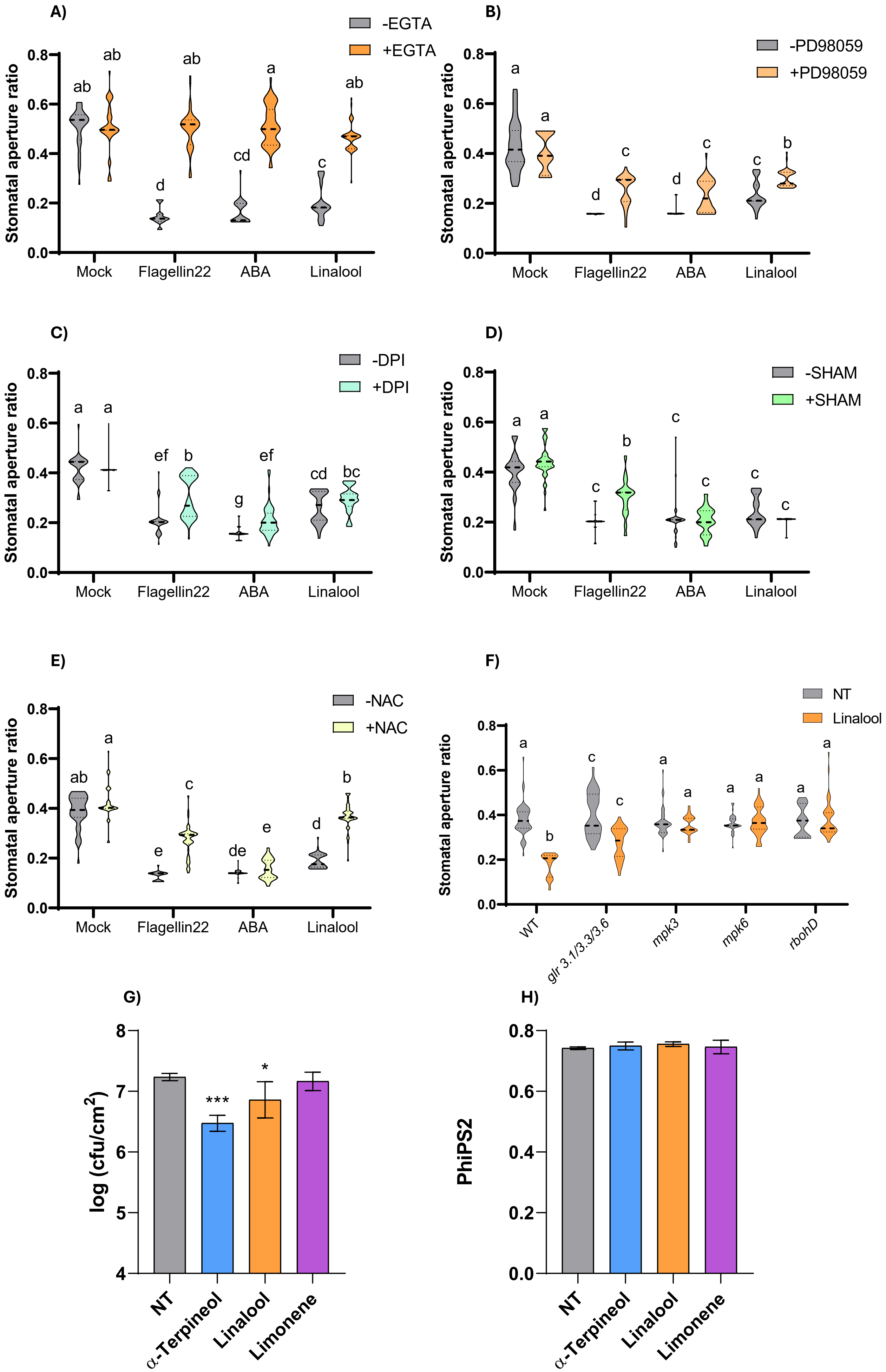


**Figure S2. Pharmacological and genetic dissection of linalool-induced stomatal responses and evaluation of monoterpenoid effects on bacterial growth and photosynthetic efficiency.** Stomatal aperture ratio was measured in epidermal peels treated with mock solution, flagelin22, abscisic acid (ABA), or linalool, in the **(A)** absence (−EGTA) or presence (+EGTA) of the calcium chelator EGTA, **(B)** absence (−PD98059) or presence (+PD98059) of the MPK pathway inhibitor PD98059, **(C)** absence (−DPI) or presence (+DPI) of the NADPH oxidase inhibitor diphenyleneiodonium (DPI) **(D)** absence (−SHAM) or presence (+SHAM) of salicylhydroxamic acid (SHAM) and **(E)** absence (−NAC) or presence (+NAC) of the antioxidant N-acetylcysteine (NAC). **(F)** Stomatal aperture ratio (width/length) measured in wild-type (WT), *glr 3.1/3.3/3.6*, *mpk3*, *mpk6*, *rbohD,* and plants under non-treated (NT) conditions or after treatment with linalool. Violin plots show the distribution of stomatal aperture ratios, with central values indicated for each treatment. Different letters indicate statistically significant differences among treatments ($p$ < 0.05). **(G)** Bacterial population size expressed as log(cfu/cm$^2$) following *Pseudomonas syringae* (Pst) infection in non treated plants (NT), monoterpenoid-treated plants (α-terpineol and linalool) and the monoterpene limonene. **(H)** Maximum quantum efficiency of photosystem II (ΦPSII) measured in non treated plants (NT), monoterpenoid-treated plants (α-terpineol and linalool) and the monoterpene limonene

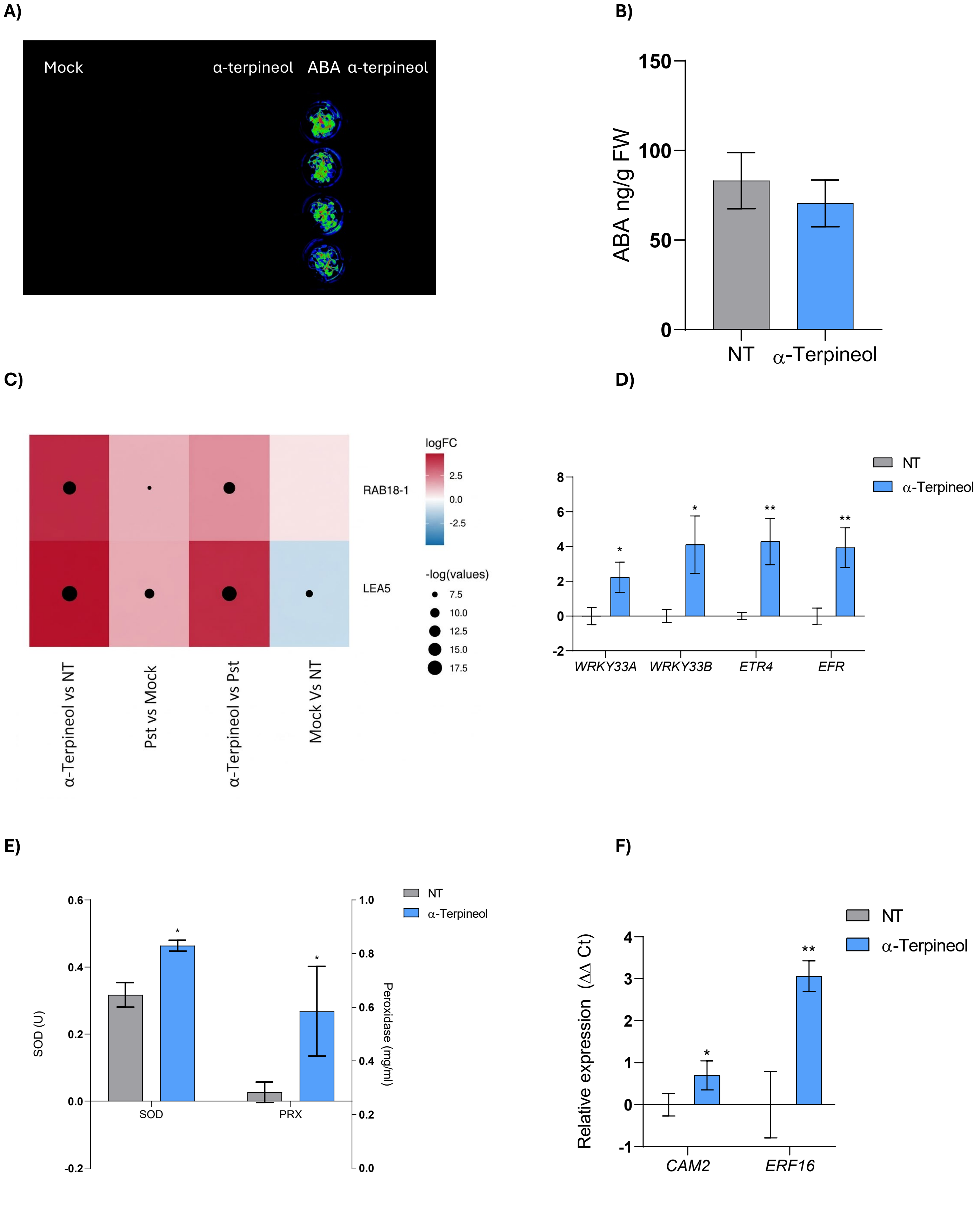


**Figure S3. Analysis of ABA-related and defence signalling responses induced by α-terpineol treatment.** **(A)** Activation of the ABA-responsive *Arabidopsis* reporter *pMAPKKK18-LUC+* following treatment with ABA or α-terpineol. **(B)** Endogenous ABA levels in non-treated (NT) and α-terpineol-treated tomato plants, showing no significant differences between treatments. **(C)** Heatmap showing the log fold change (logFC) of selected ABA-related marker genes (*RAB18-1* and *LEA5*) in α-terpineol-treated plants and Pst-infected plants relative to their respective controls. Dot size represents the −log(*p*-value) of differential expression. **(D)** Relative expression of MAPK-associated genes (*WRKY33A*, *WRKY33B*, *ETR4*, and *EFR*) in tomato leaves treated with α-terpineol compared to NT controls. **(E)** Activity of ROS-detoxifying enzymes superoxide dismutase (SOD) and peroxidase (PRX), in response to α-terpineol treatment. **(F)** Relative expression of $Ca^{2+}$-responsive genes (*CAM2* and *ERF16*) in tomato leaves treated with α-terpineol. Gene expression levels were normalized to reference genes and are presented as relative expression values (ΔΔCt). Data represent mean ± SE of at least three independent biological replicates. Asterisks indicate statistically significant differences compared with NT controls (* $p < 0.05$; ** $p < 0.01$).

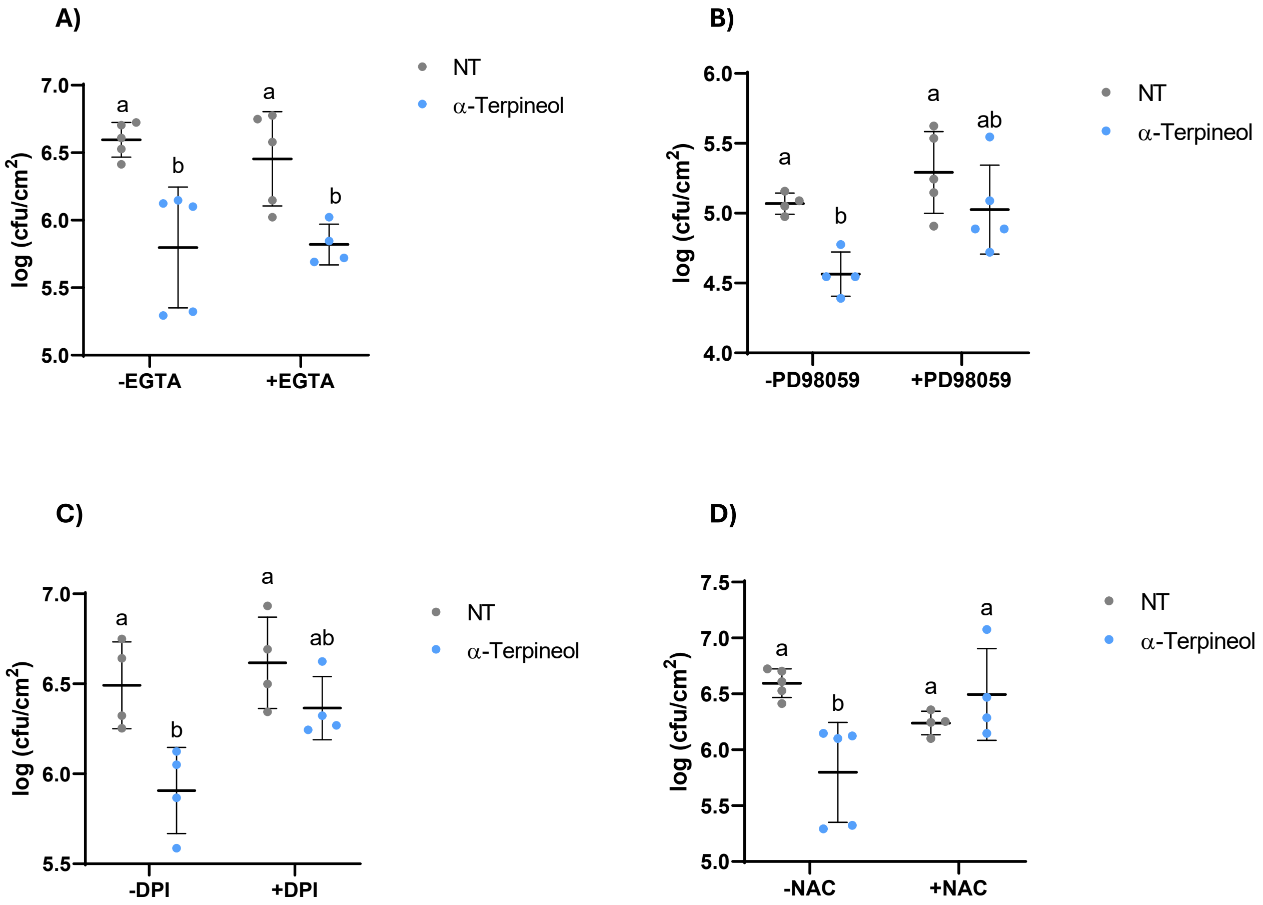


**Figure S4. Effect of pharmacological inhibitors on α-terpineol-mediated resistance to *Pseudomonas syringae*.** Bacterial growth is expressed as log of colony-forming units per $cm^2$ log($cfu/cm^2$) in non-treated (NT) and α-terpineol–treated plants in the **(A)** absence (−EGTA) or presence (+EGTA) of the calcium chelator ethylene glycol tetraacetic acid (EGTA), **(B)** absence (−PD98059) or presence (+PD98059) of the MAP kinase pathway inhibitor PD98059, **(C)** absence (−DPI) or presence (+DPI) of the NADPH oxidase inhibitor diphenyleneiodonium (DPI), **(D)** absence (−NAC) or presence (+NAC) of the antioxidant N-acetylcysteine. Each point represents an individual biological replicate; horizontal bars indicate mean values ± SE. Different letters indicate statistically significant differences among treatments ($p < 0.05$).

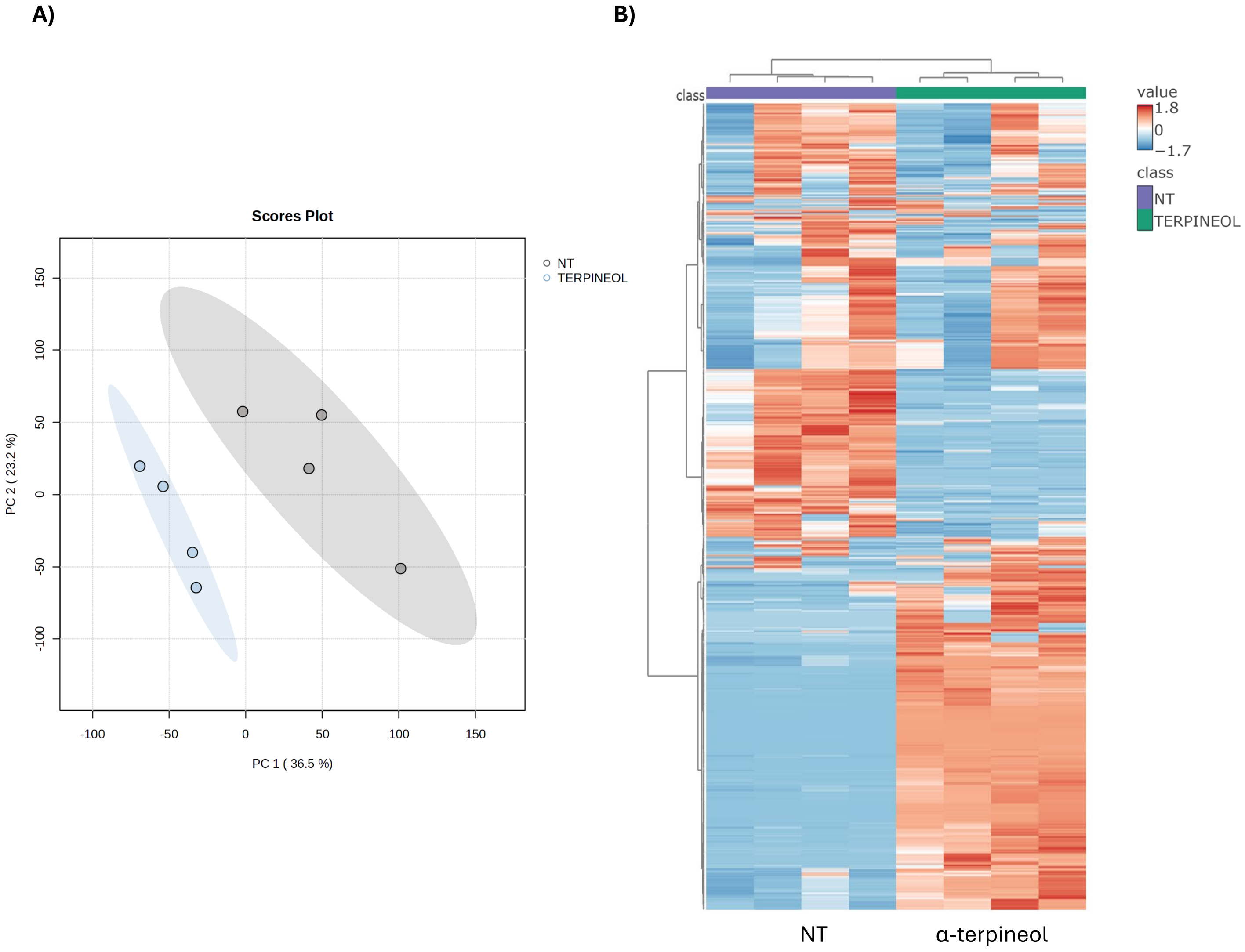


**Figure S5. Multivariate and compositional analysis of volatile profiles following α-terpineol treatment. (A)** Principal component analysis (PCA) scores plot showing the separation between non-treated (NT) and α-terpineol–treated samples based on volatile compound profiles. Each point represents an individual biological replicate, and shaded areas indicate group clustering. **(B)** Heatmap representation of volatile compounds detected in non-treated and α-terpineol–treated samples. Rows correspond to individual volatile features and columns to biological replicates. Colour scale indicates relative abundance values, with hierarchical clustering applied to both samples and compounds.

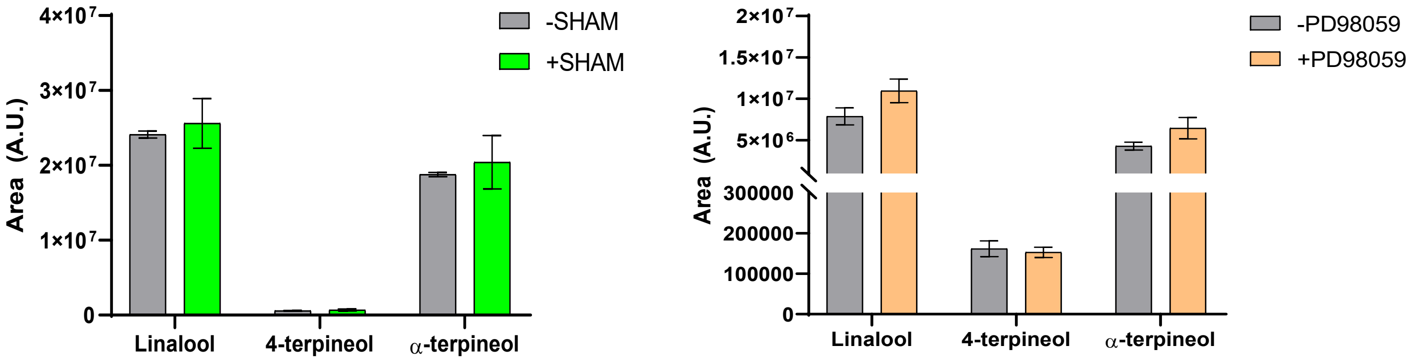


**Figure S6. Effect of SHAM and PD98059 treatments on HMTP levels during *Pseudomonas syringae* pv. *tomato* infection.** Accumulation of linalool, α-terpineol, and 4-terpineol was quantified as peak area (arbitrary units, A.U.). Prior to infection with *Pseudomonas syringae* pv. *tomato*, *in planta* treatments with the indicated chemical inhibitors (SHAM or PD98059) were performed, with corresponding untreated RG tomato plants used as controls. Data represent mean ± standard error. No statistical differences were detected.

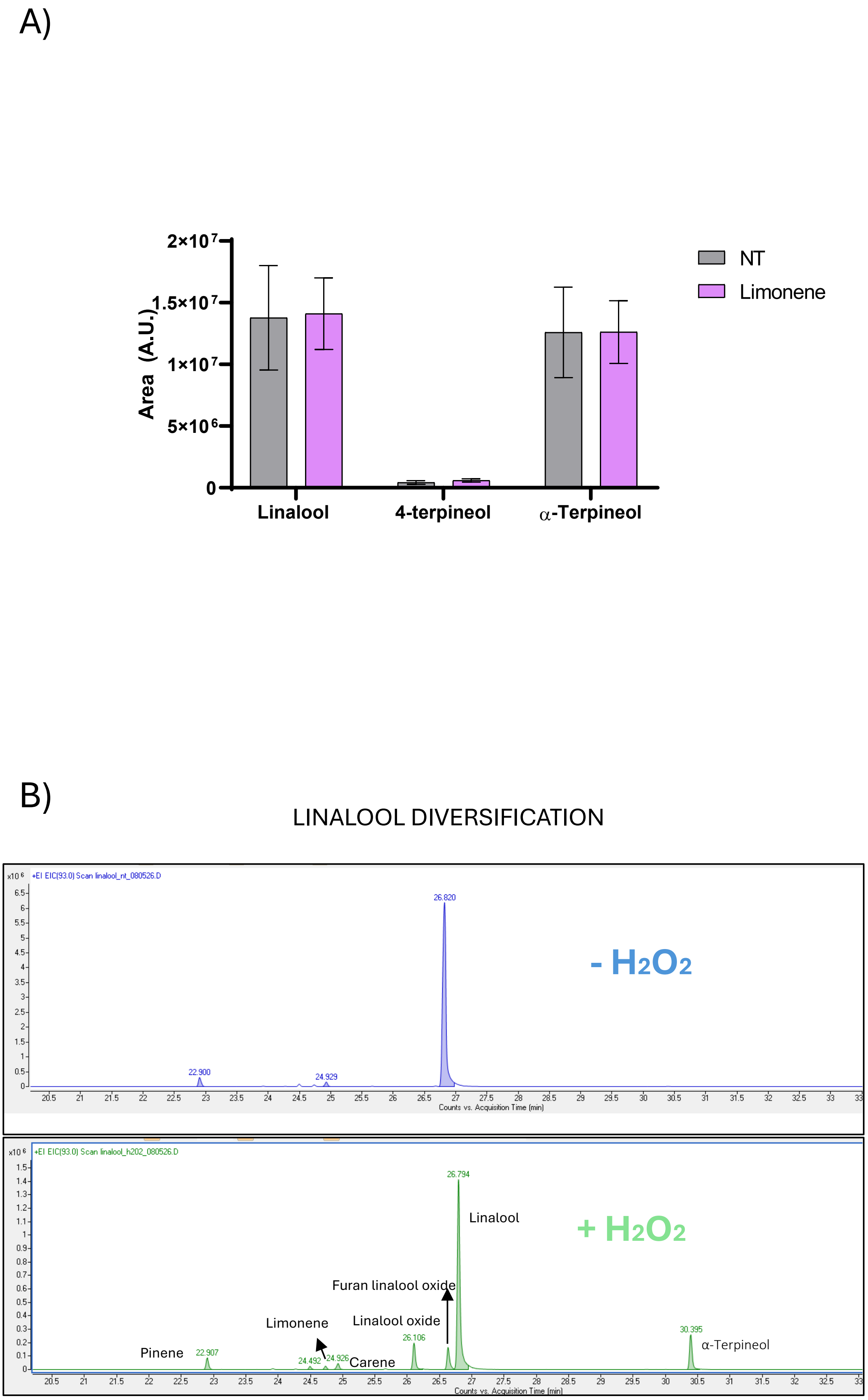


**Figure S7. Linalool is converted into other monoterpenes under oxidative stress while limonene treatments do not change monoterpenoid content. (A)** Relative abundance (area arbitrary units, A.U.) of linalool, 4-terpineol, and α-terpineol in non-treated (NT) samples and samples treated with limonene. Error bars represent standard deviation Error bars represent standard deviation. **(B)** Extracted ion chromatograms (EICs) of linalool (Retention Time, RT 26,794) analysed in the absence ($-H_2O_2$, top panel, blue) and presence ($+H_2O_2$, bottom panel, green) of hydrogen peroxide.

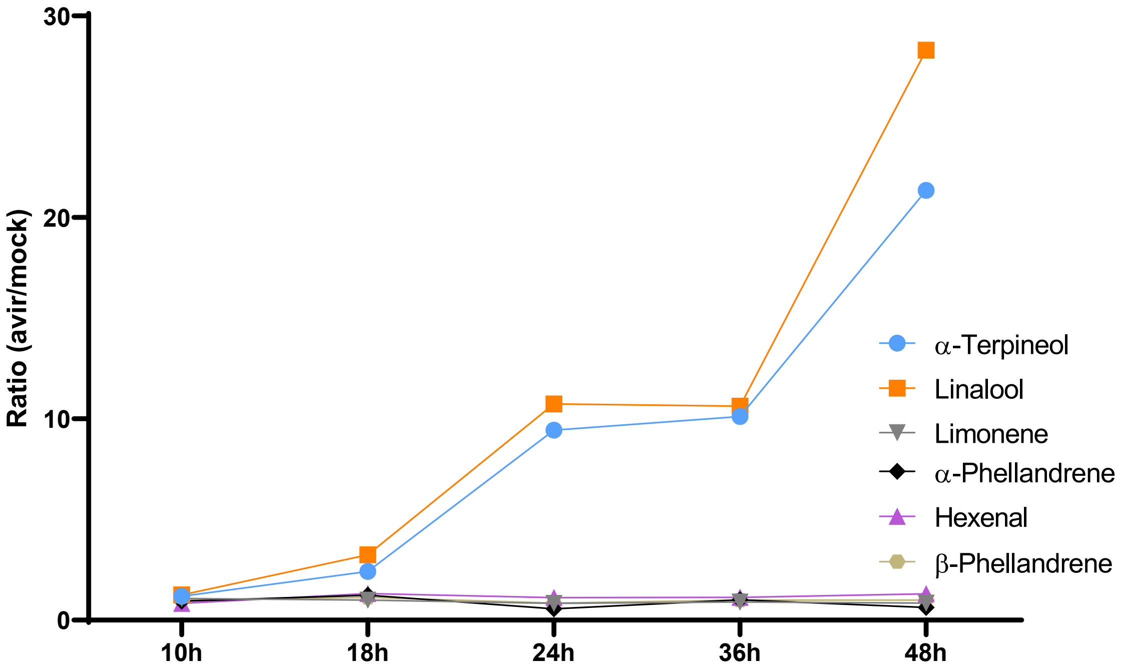


**Figure S8. Early temporal dynamics of volatile accumulation during avirulent *Pseudomonas syringae* infection.** Relative accumulation of hydroxylated monoterpenes (α-terpineol and linalool) and the non-hydroxylated monoterpenes (hexenal, limonene, α-phellandrene and β-phellandrene) at early time points (10, 18, 24, 36, and 48 h) following avirulent *Pseudomonas syringae* infection. Data are expressed as ratios between infected and mock-treated plants.

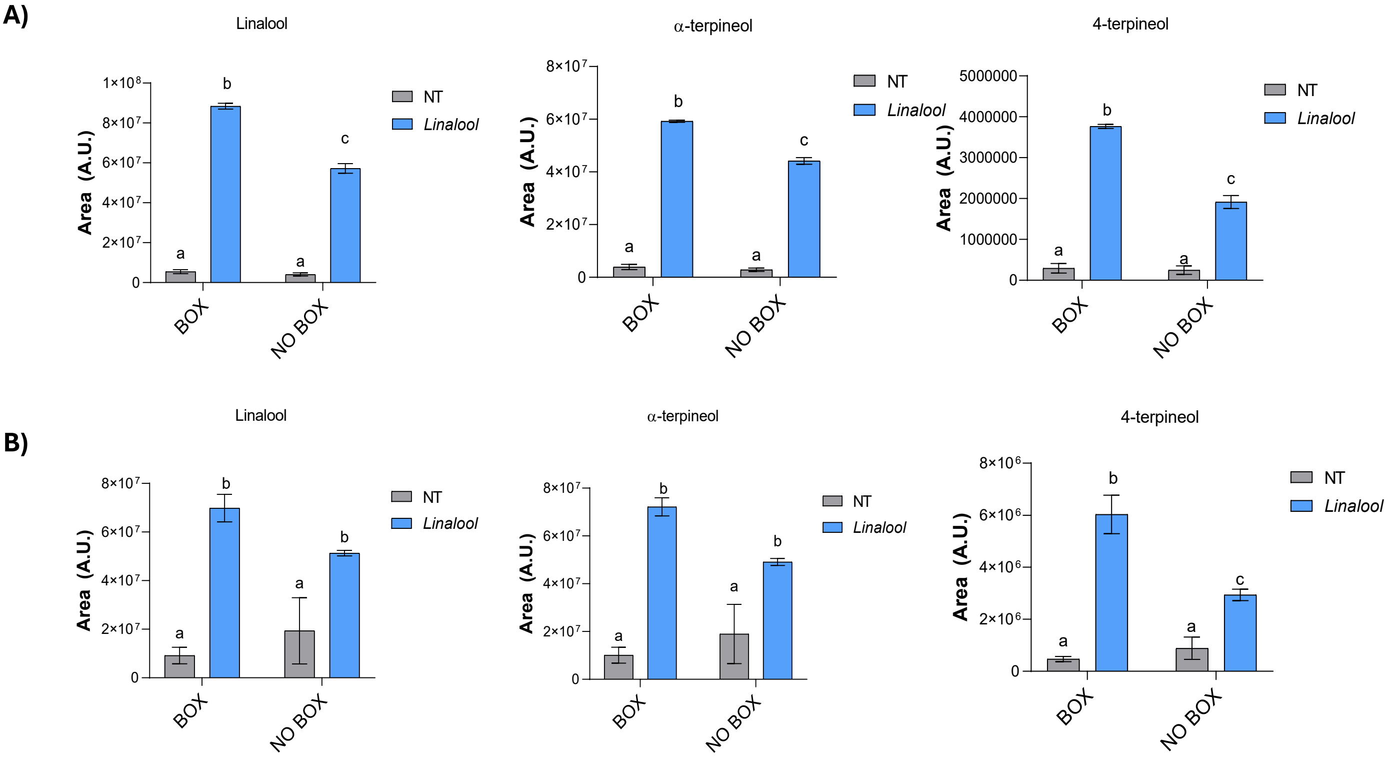


**Figure S9. HMTP accumulation in plants maintained inside or outside closed boxes.** Levels of linalool, α-terpineol, and 4-terpineol were measured in tomato plants treated with linalool and maintained either inside closed boxes (BOX) or outside the boxes (NO BOX) for 24 h **(A)** or 72 h **(B)**. Volatile compounds were quantified by GC–MS, and are expressed as peak areas (arbitrary units, A.U.). NT, non-treated plants. NT, non-treated plants. Italicized "linalool" refers to the treatment applied to the plants, whereas non-italicized compound names indicate the detected volatile levels. Bars represent mean values ± SD. Two-way ANOVA was performed. Different letters indicate statistically significant differences ($p < 0.05$).

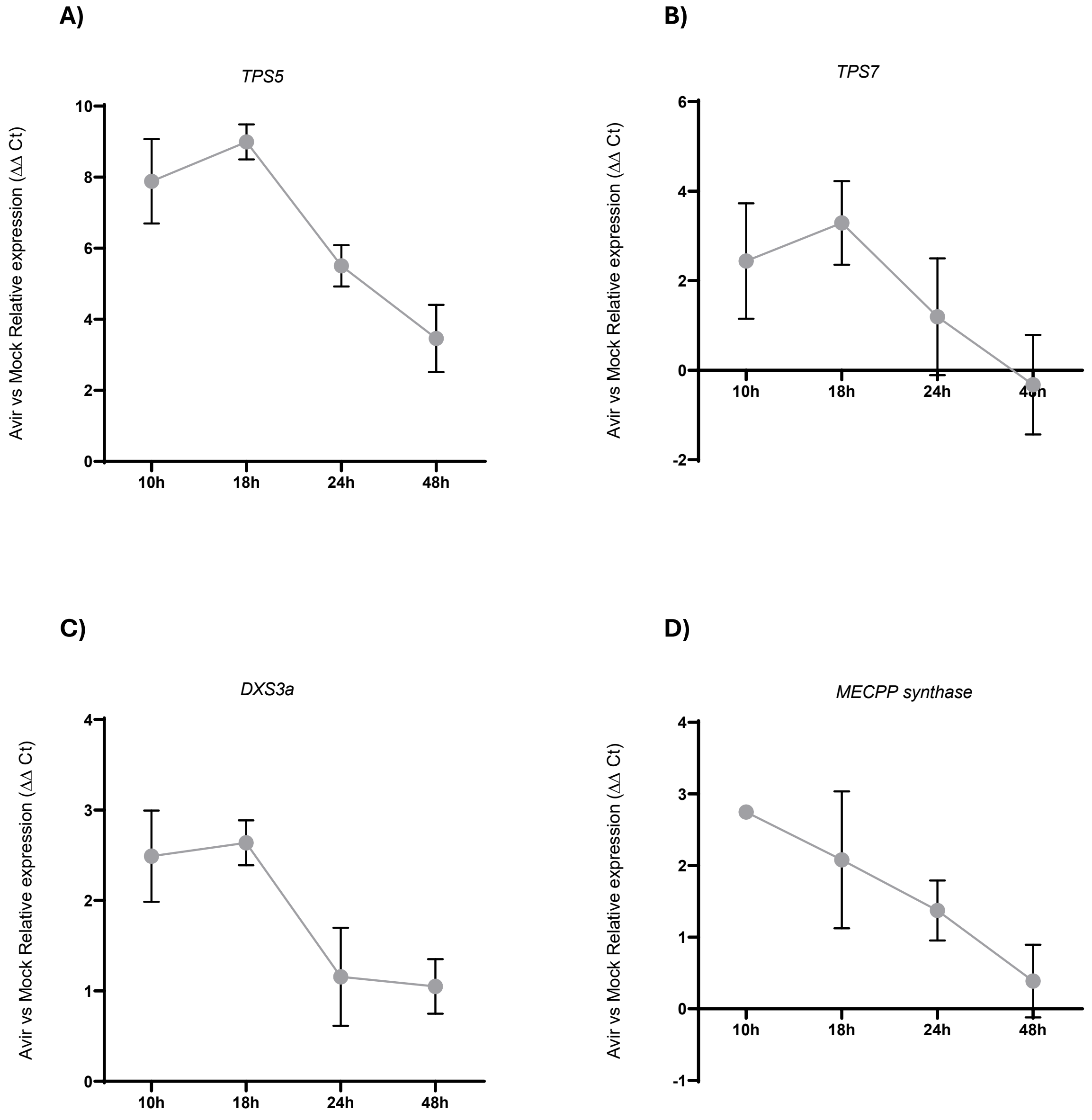


**Figure S10. Temporal dynamics of terpene biosynthesis gene expression during avirulent *Pseudomonas syringae* infection.** Relative expression (ΔΔCt, Avir vs. Mock) of terpene synthase genes *TPS5* (linalool-associated) **(A)** and *TPS7* (limonene-associated) **(B)**, as well as MEP pathway genes *DXS3a* **(C)** and *MECPP synthase* **(D)**, are shown at 10, 18, 24, and 48 hours post-inoculation. Data are expressed as ratios between infected and mock-treated plants.

| Gene | Forward | Reverse |
|---|---|---|
| *PR1* | *5' ACT CAA GTA GTC TGG CGC AAC TCA 3'* | *5' AGT AAG GAC GTT GTC CGA TCG AGT 3'* |
| *Actin* | *5' CTA GGC TGG GTT CGC AGG AGA TGA TGC 3'* | *5' GTC TTT TTG ACC CAT ACC CAC CAT CAC AC 3'* |
| *WRKY33A* | *5' GCA TTA CTG TCA ACC ATC GC 3'* | *5' AAC TTC GCG GAT TCT CAC TT 3'* |
| *WRKY33 B* | *5' CCA CAA CAG TCT GAA ATG GG 3'* | *5' CAG CAA AGC AAT GAC TCC AT 3'* |
| *MEcPP Synthase* | *5' TGG CGT CTA GGT TTC CCA AT 3'* | *5' CAA TTT TGG GAG CTC TGG GG 3'* |
| *DXS3a* | *5' GTC CAC AGC AAG CAA AAG GA 3'* | *5' GCT TCT ACA CAT GCC TGC TC 3'* |
| *TPS7* | *5' ATG CCC ATC TCC ATG CTG AT 3'* | *5' CAT GAA TGA CAC GGG TGC TT 3'* |
| *TPS5* | *5' TGG TGG TCA CCT TCA AGA GA 3'* | *5' GCC TTG TGG AAA TAG GA 3'* |
| *MYC2* | *5' GCT TCC AGT GCC AAT GTG AA 3'* | *5' CCC GAA GAA GGC AAA ACT GT 3'* |
| *WRKY 75* | *5' TTT CCA TCA GCA TCA TCG TC 3'* | *5' GGC CTT ATT CTT CCC TTG GA 3'* |
| *GA200X* | *5' CTC ATT TCT AAT GCT CAT CGT 3'* | *5' TGA GAT GAT TCT TTC TTA GCG 3'* |
| *NAC* | *5' GGA AGG CTA CTG GCA CTG AT 3'* | *5' TGA CAC CTT TTG GTG GTT TG 3'* |
| *CAM2* | *5' AGG AGG AGT TGA AAG AGG CAT TC 3'* | *5' CTC TCC AAG GTT AGT CAT CAC ATG A 3'* |
| *P23/PR5* | *5' TTC GAG` GTA CGC AAC TG 3'* | *5' TGC ATT GAT GAC CCA TGT TT 3* |

**Supplemental Table S1.** Primer sequences used for RT-qPCR analyses.
